# Nucleo-cytoplasmic trafficking regulates nuclear surface area during nuclear organogenesis

**DOI:** 10.1101/326140

**Authors:** Vincent Boudreau, James Hazel, Jake Sellinger, Pan Chen, Kathryn Manakova, Rochelle Radzyminski, Hernan Garcia, Jun Allard, Jesse Gatlin, Paul Maddox

**Affiliations:** Department of Biology, University of North Carolina at Chapel Hill; Physiology Course, Marine Biological Laboratory; Department of Molecular Biology, University of Wyoming; Center for Complex Biological Systems, University of California, Irvine; Department of Physics, University of California, Irvine; Department of Mathematics, University of California, Irvine; Department of Physics, University of California at Berkeley, Berkeley, California; Department of Molecular & Cell Biology,University of California at Berkeley, Berkeley, California; Institute for Quantitative Biosciences-QB3, University of California at Berkeley, Berkeley, California; Biophysics Graduate Group, University of California at Berkeley, Berkeley, California

## Abstract

Throughout development, nuclei must be assembled following every cell division to establish a functional organelle from compact, mitotic chromatin. During nuclear organogenesis, chromatin expands to establish a nucleus of a given size seperate from the cytoplasm. Determining how nuclear organogenesis is regulated is particularly significant in the context of certain cancers in which scaling relationships between cell and nuclear sizes are not maintained. Controlling cell size in vitro using a microfluidics approach, we determined that neither nuclear volume nor surface area scale directly with cell size. Looking to explain differential nuclear scaling relationships, we developed a simple mechano-chemical mathematical model. In simulating biological perturbations in silico, our model predicted crucial roles for nucleo-cytoplasmic trafficking in regulating nuclear expansion and in restricting the recruitment of a potential nuclear surface area factor. In mammalian tissue culture, inhibiting nuclear export increased nuclear expansion rates and reduced the amount of nuclear lamin, a candidate surface area factor, being recruited to assembling nuclei, supporting our model’s predictions. Targeting the principal nuclear export component in the Drosophila syncytial embryo, Embargoed, we show that nuclear expansion rates are also increased in this developmental context, consistent with our model. Using the MS2-reporter system in fly embryos, we demonstrate a role for nuclear export in regulating transcription activation timing and dynamics, suggesting that regulating nuclear assembly is crucial for downstream nuclear function. Taken together, we propose a simple model through which nuclear organogenesis is achieved and demonstrate a role for nuclear export in regulating nuclear assembly.

## INTRODUCTION

Assembling the nucleus into a functional organelle is a regulated, stepwise process essential for nuclear function. Most of our understanding of nuclear organogenesis stems from the examination of nuclear envelope reformation (NERF) at the end of mitosis in organisms that undergo nuclear envelope breakdown (NEBD) at mitotic entry (reviewed in Güttinger et al., 2009). Although the temporal recruitment of many nuclear assembly components during mitotic exit and their regulation has become clearer, how the nucleus expands into a functional organelle within a cell of a limited volume remains elusive.

NERF is a stepwise process implicating several biochemical cues that sequentially trigger chromosome arm compaction (Mora-Bermudez et al., 2007), the recruitment of nuclear pore (Mansfeld et al., 2006; Rasala et al., 2006; Stavru et al., 2006) and membrane precursors (Hallberg et al., 1993), and nuclear lamina assembly (Tsenga and Chena, 2011). PP2A-B55, the principal CyclinB-CDK1 substrate dephosphorylating component during mitotic exit (Mochida et al., 2009; Gharbi-Ayachi et al., 2010), is one of the core biochemical factors driving nuclear expansion and NERF, and is known to collaborate with Importin β (Impβ, Schmitz et al., 2010). The involvement of Impβ, and of nucleo-cytoplasmic trafficking (NCT) generally, in regulating nuclear assembly has long been suggested by several lines of evidence, including the requirement of Ran’s GTPase activity in NERF (Hetzer et al., 2000). Additionally, the nuclear export factor Crm1 has been shown to be important for chromosome segregation (Arnaoutov et al., 2005) and the translocation of the chromosomal passenger complex from the centromere to the midzone in anaphase (Knauer et al., 2006). Therefore substantial evidence suggests that both protein import and export pathways are important in mitotic exit models of nuclear organogenesis. Despite the highlighted role for NCT, there is little mechanistic information to suggest how NCT regulates nuclear organogenesis or what molecules must be trafficked. It also remains unclear how disrupting NCT during nuclear assembly affects function in the subsequent interphase.

Another important question in understanding nuclear organogenesis is how nuclei expand to a given nuclear size in a given cellular volume. It has been well established that cell size (or cytoplasmic volume) scales with nuclear size in different metabolic and genetic conditions (Jorgensen et al., 2007; Neumann and Nurse, 2007) and in different developmental situations (Levy et al., 2010). The mechanisms by which nuclear size is regulated, particularly in the context of genomic content, are less clear as genetic perturbations modifying ploidy by an order of magnitude have been reported to have minimal effects on nuclear size (Neumann and Nurse, 2007). In contrast, the increase in genomic content following DNA replication has been reported to cause a 20% to 100% increase in nuclear volume (Jorgensen et al., 2007), whereas inhibiting replication has been shown to result in a significant decrease in nuclear size (Levy et al., 2010). In a model in which nuclear size is dependent on genomic content, genomic content and nuclear volume should increase linearly with one another whereas nuclear volume should be unaffected by changes in cytoplasmic volume (Fig. 1A). An alternative nuclear scaling model in which nuclear size is determined by nucleoplasmic or nuclear envelope components (Fig. 1B) would be independent of genomic content. This model would be in line with current evidence of nuclear to cell size scaling (Jorgensen et al., 2007, Levy et al., 2010, Neumann and Nurse, 2007), yet it remains unclear whether nucleoplasmic or nuclear envelope components set nuclear size.

**Figure 1.**
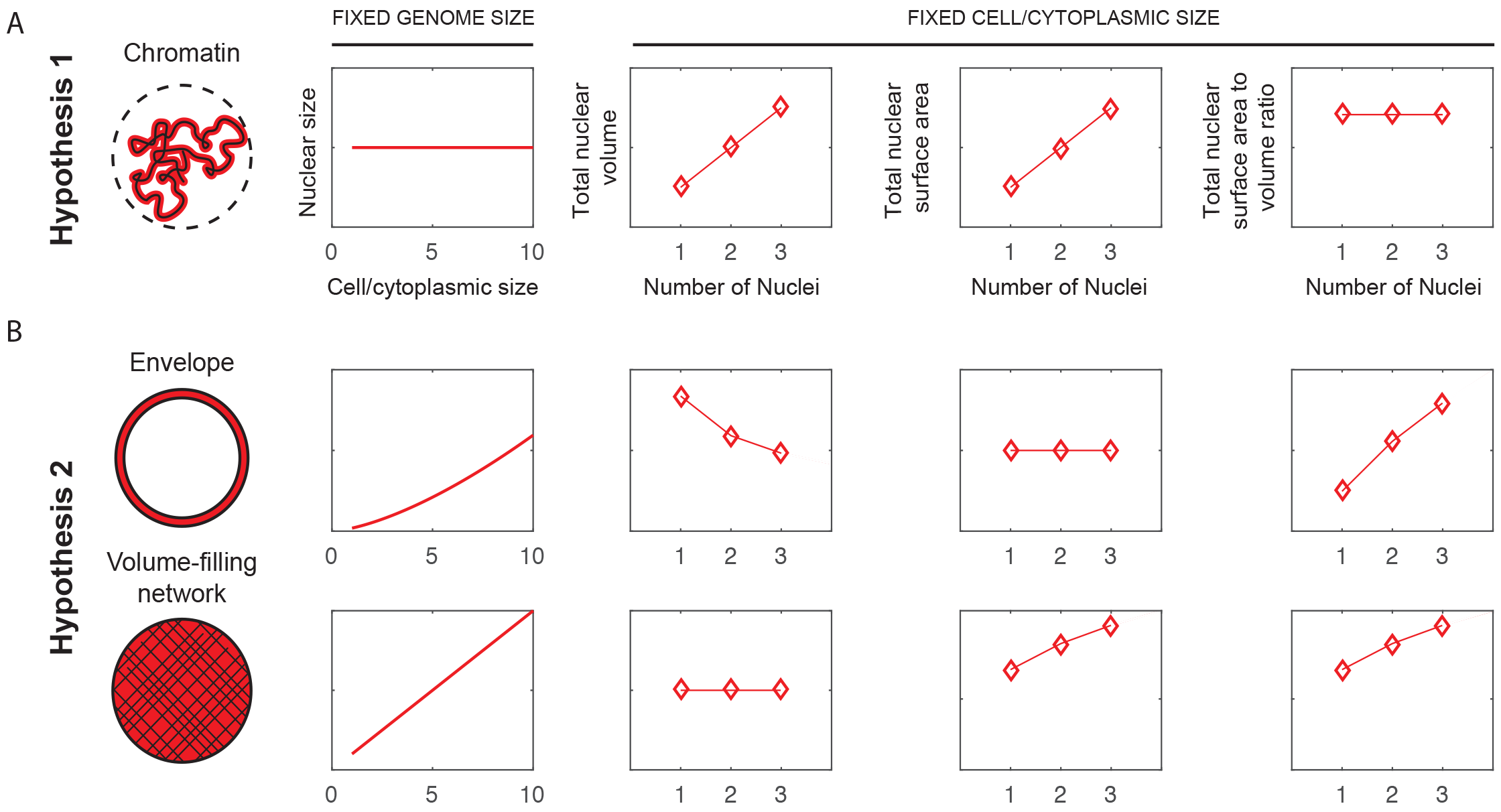
Models of nuclear size scaling with cell size and nuclear surface area to volume ratios within a defined cell size. Simple models of nuclear size in which either the number of nuclei/genome size or cell size/cytoplasmic size are fixed predict nuclear size scaling relationships and nuclear surface area:volume ratios under different regimes. **a,** In Hypothesis 1, nuclear size is driven by genome size. As cell size/cytoplasmic size increases, nuclear size is constant as genome size is constant. By keeping cell size constant and varying the number of genomes in this regime, nuclear volume and nuclear surface area increase. **b,** In Hypothesis 2, nuclear size to cell size scaling is independent of genome size. Considering the nucleus as a viscoelastic material composed of a nuclear envelope and a nucleoplasmic volumetric network, nuclear size is hypothesized to be regulated in a nuclear envelope-limited regime (upper panel) or a volumetric network-limited regime (lower panel), where either nuclear surface or nuclear volume set nuclear size respectively. * = p < 0.05.

To examine how nuclear organogenesis is regulated by genomic content, nuclear volumetric and surface area components specifically and NCT broadly, we examined nuclear volume and nuclear surface area scaling relationships, developed a mathematical model and tested its predictions in cells and in a developing organism. First, in gaining biophysical control over cell size, we measured nuclear volume and nuclear surface area *in vitro* while varying the number of nuclei in a given cytoplasmic volume using a previously described microfluidics approach in X. laevis egg extracts (Hazel et al., 2013). Second, using measured concentrations of proposed volumetric and surface area regulating factors (Wühr et al., 2014) as well as nuclear expansion measurements in cytoplasmic volume-restricted conditions, nuclear assembly was modelled. Taking advantage of this model trained solely on *in vitro* data from Xenopus egg extracts, we predicted *in silico* and tested in cells and in vivo how NCT regulates nuclear organogenesis. These results uncover a role for nuclear import and nuclear export in regulating nuclear expansion.

Nuclear export was predicted and found to regulate nuclear expansion through the localization of a proposed surface area factor (Lamin B). In developing *D. melanogaster* embryos, compromising NCT affects nuclear expansion in a manner consistent with our model. To test how NCT’s role in nuclear organogenesis affects downstream nuclear function, we show that disrupting NCT impairs transcription activation in interphase following nuclear assembly using an *in vivo* live-cell transcription reporter (Garcia et al., 2013). Overall, we demonstrate NCT regulates nuclear expansion through proposed nuclear surface area factors during nuclear organogenesis - a process required for downstream nuclear function.

## RESULTS

### Neither nuclear surface area nor nuclear volume scale with cell size in multinucleate cytoplasm droplets

Nuclear, organelle, chromosome, and spindle size have all been shown to scale with cell size in either developmental or in vitro contexts (Hazel et al., 2013; Ladouceur et al., 2015; Good et al., 2013; Wilbur et al., 2013). How the contribution of genomic content and cytoplasmic volume define a given nuclear size have not been examined with controlled biophysical variables.

Here, a previously described microfluidics approach (Hazel et al., 2013) was used to encapsulate fully functional cytoplasm (derived from X. laevis eggs; Desai et al., 1999) of defined volumes to probe how physical parameters affect nuclear assembly processes. Using de-mem-branated sperm nuclei as a source of chromatin in addition to 2 μM of GF-P:NLS, droplets containing mainly spherical nuclei were observed, which expand and reach steady state within the duration of the experiment.

To determine how genomic content, nuclear volume, and nuclear surface area influence nuclear assembly, we measured nuclear size in cytoplasmic droplets of fixed volumes (50μm in diameter) containing a single or several genomes (Fig. 2A). Whereas total nuclear volume increased slightly in droplets containing two or more nuclei (Fig. 2B), total nuclear surface area increased faster (Fig. 2C). This observation is inconsistent with models in which nuclear size is driven solely by chromatin amount (Fig. 1A). Additionally, total nuclear volume and surface area both increased in droplets with multiple nuclei, indicating that nuclear size is likely not dictated by nucleoplasmic or nuclear envelope components exclusively, as hypothesized in Figure 1B. Indeed, these data suggest nuclear size is regulated by a combination of nuclear envelope and nucleoplasmic components, in addition to chromatin amount.

**Figure 2.**
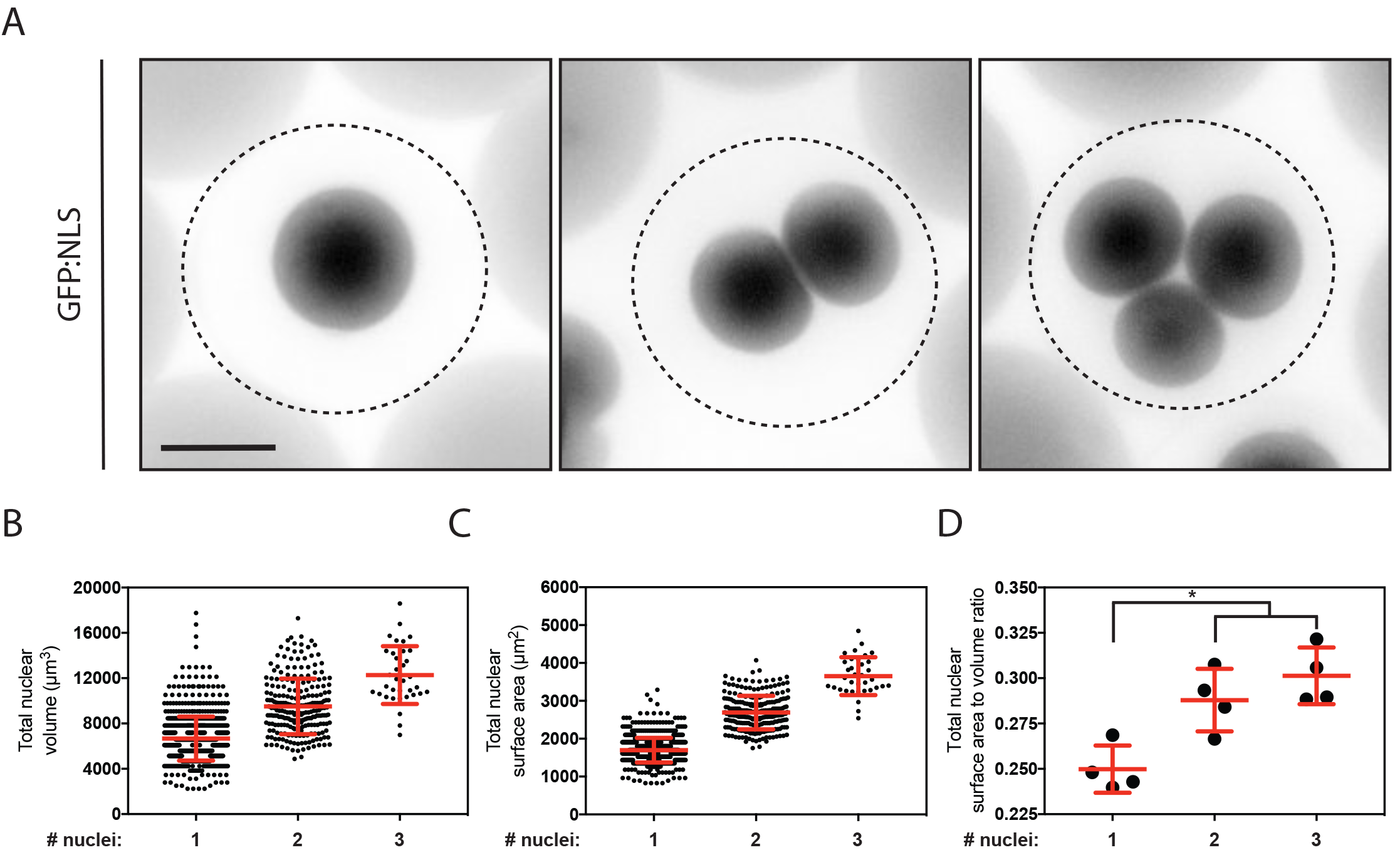
in vitro encapsulation of nuclei in cytoplasmic droplets reveals nuclear surface area to volume regulation during nuclear expansion. **a,** Representative images of 1, 2, or 3 nuclei assembled in droplets with a diameter of 50 μm. Nuclei were visualized by import of an GFP:NLS reporter to assess nuclear morphology. Scale bar, 20 μm. **b,** Total nuclear volume plotted as a function of the number of encapsulated nuclei at 90 minutes post encapsulation in droplets (n > 1000). **c,** Total nuclear surface area plotted as a function of the number of encapsulated nuclei at 90 minutes post encapsulation (n > 1000). **d,** Surface area to volume ratios of mean surface areas and volumes of nuclei 90 minutes post encapsulation in droplets across four experimental replicates. Error bars represent SD.

### Modelling nuclear organogenesis captures the roles of nuclear volumetric and surface area factors in regulating nuclear expansion

To explore conditions under which nuclear scaling would be expected to occur, we developed a mathematical model of the mechanochemistry of nuclear assembly. In the model, the nucleus is comprised of a viscoelastic body (containing chromatin and internal architectural networks such as NuMA that provide mechanical rigidity) surrounded by a viscoelastic envelope (the lamina and membranes). The fundamental assumption of the model is that two molecular components must be imported into the nucleus for its growth: a surface factor that assembles into the envelope and a volume factor that assembles into the interior. We do not a priori specify the molecular identity of these components.

Nuclear size is assumed to change in response to hydrostatic pressure and surface tension (Kim et al., 2016). Surface tension is due to a surplus or deficit of surface factor. Hydrostatic pressure is due to a surplus or deficit of volume factor as well as the confinement of chromatin. The model assumes both the surface factor and volume factor are uniformly distributed throughout the cytoplasm and must be transported by passive diffusion and microtubule-associated dynein to the nuclear periphery. Once the factor is near the nucleus, it is imported to (and exported from) the nucleus, enabling nuclear growth. A full mathematical description is given in Supplemental Math.

We use this model to simulate nuclear expansion following the anaphase-telophase transition. For a given set of biophysical input parameters (nuclear rigidity, import and export rates), the simulation produces time trajectories of nuclear size (Fig. 3B) and the concentrations of volume and surface factors. The model captures experimental nuclear expansion measurements of expanding nuclei in cytoplasmic droplets (Fig. 3C and 3D), in which nuclear size reaches steady-state over the course of the experiment (Fig. 3C).

Under Hypothesis 2 (Fig 1B), the model demonstrates that, depending on the ratio of nuclear surface rigidity to nuclear volume factor rigidity, multi-nucleated droplets could scale with either constant total volume or constant total surface area (SuppMath Fig. S3), or combinations of these. Notably, all Hypothesis 2 simulations give non-increasing total nuclear volumes. Our experimental data (Fig. 2 and Fig. 3) place the model parameters in a regime in which nuclear volume rigidity is dominant, or, equivalently, in the regime in which the surface factor is not limited, and that chromatin confinement contributes significantly to nuclear size (SuppMath Fig. S4), which we refer to as Hypothesis 3 (Fig. 3E). By changing biophysical parameters, the model can simulate perturbations including to import/export or microtubules. This is discussed below (Fig 4A, 4B and 5B).

**Figure 3.**
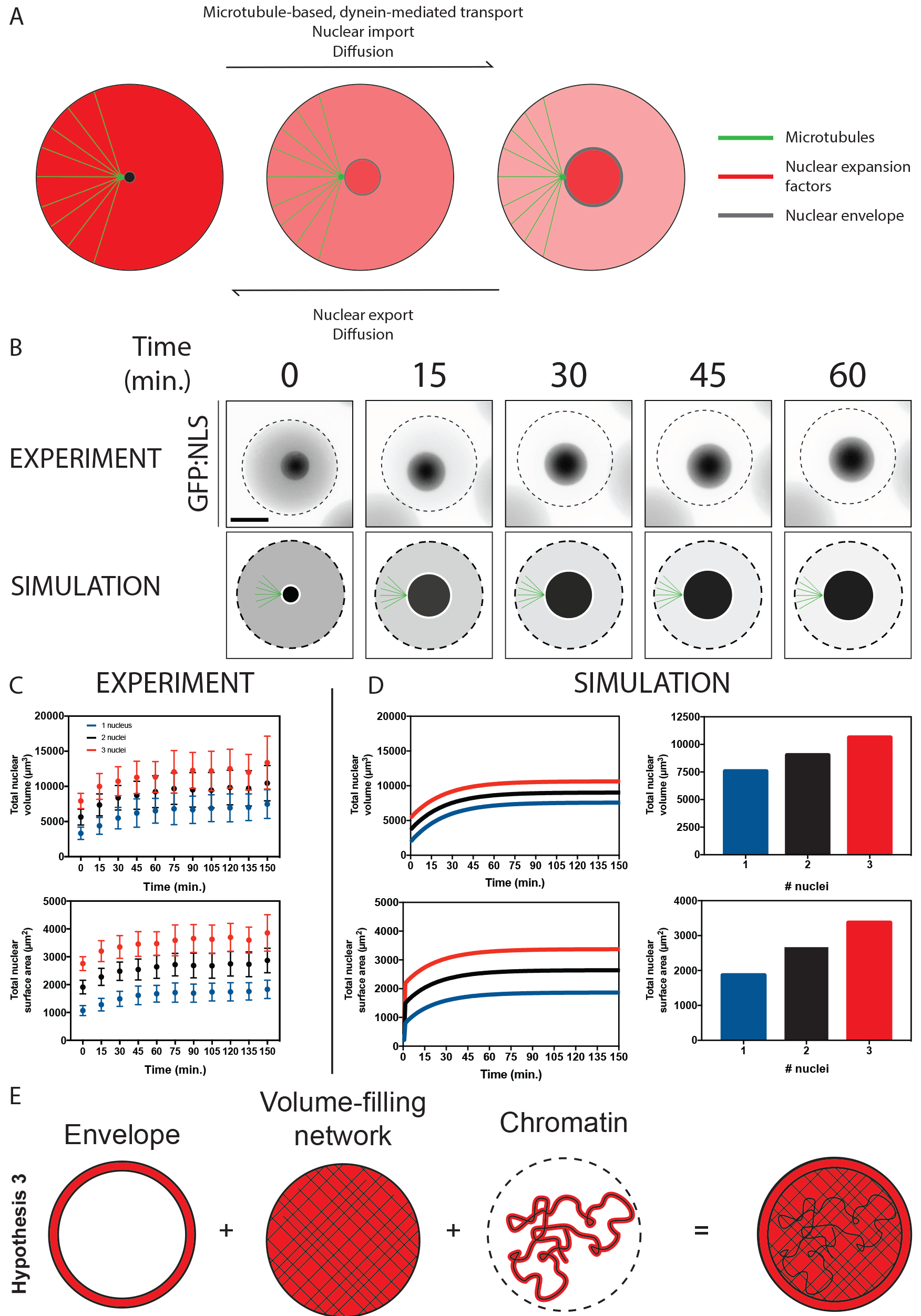
A mechano-chemical mathematical model dependent on dynein-based transport supports an inverse scaling relationship between nuclear diameter and nuclear surface area. **a,** Model assumptions of regulatory mechanisms governing nuclear surface area and volumetric factor concentration. The model assumes factors can become proximal to the expanding nucleus based on diffusion and dynein-based transport and are regulated at the nuclear envelope by nucleo-cytoplasmic trafficking. **b,** Experimental and simulated expansion of single nuclei in a defined cytoplasmic volume. Experimental nuclear expansion was visualized using a GFP:NLS reporter in encapsulated cytoplasmic droplets of ~50 μm in diameter. Dashed lines represent cytoplasm edges. **c,** Quantification of sum nuclear volume and nuclear surface area in experimental droplets containing one or several nuclei. Error bars represent SD. **d,** Total nuclear surface area and nuclear volume during simulated nuclear expansion and at nuclear expansion steady state. **e,** Quantification of nuclear expansion in defined cytoplasmic volumes suggests a model in which nuclear organogenesis occurs in a regime in which genomic size, nucleoplasmic factors and nuclear envelope factors collaborate.

**Figure 4.**
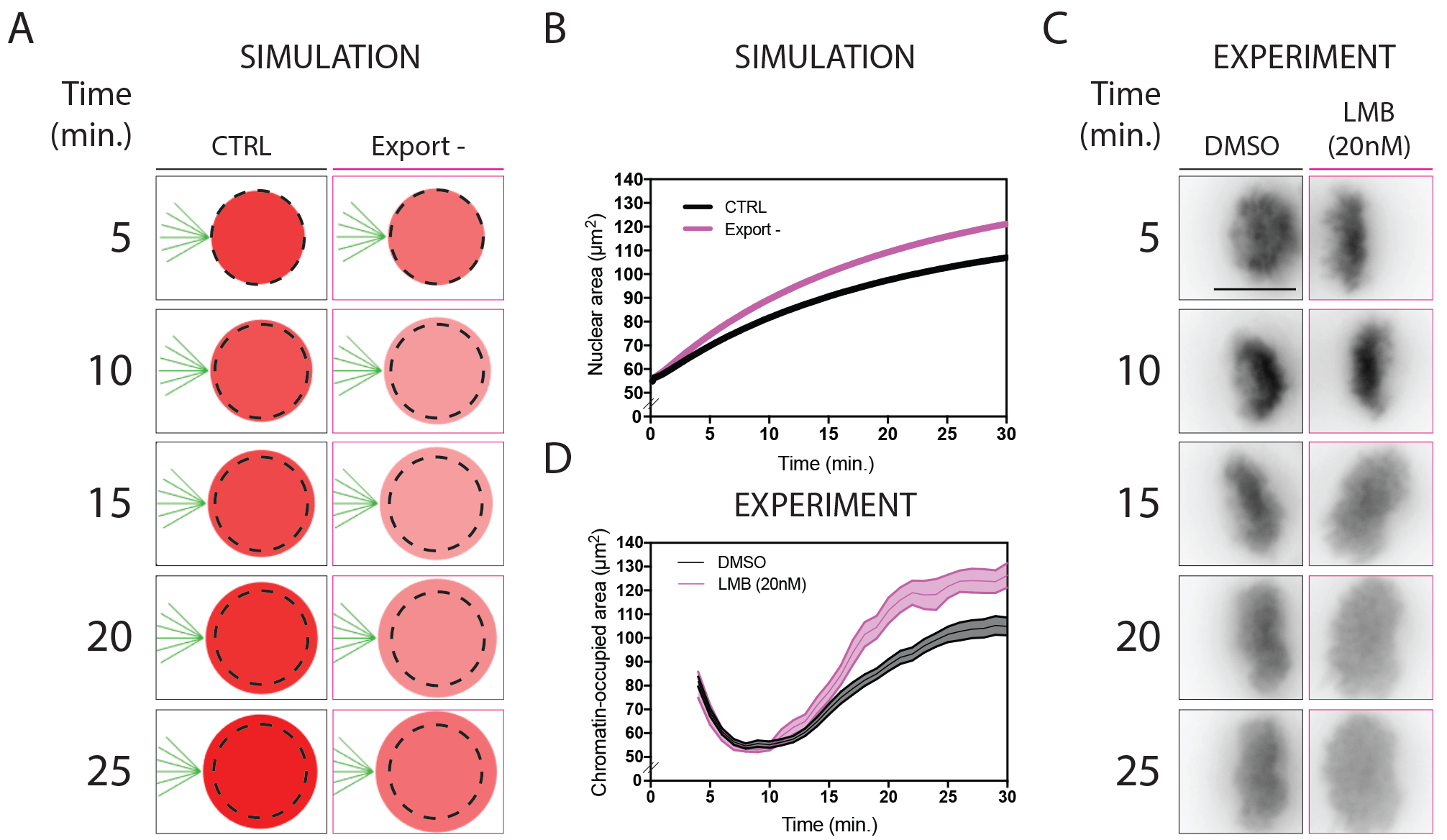
Nucleo-cytoplasmic trafficking regulates nuclear expansion. **a,** Images of simulated nuclei undergoing nuclear expansion in a defined cytoplasmic volume. Nuclear export is compromised by 50% in this simulation. Dashed lines in all images represent nuclear size at t = 5 minutes in each condition. Shades of red represent the predicted nuclear concentrations of nuclear surface area factor, **b,** Predicted nuclear cross-sectional area of simulated nuclei through nuclear expansion in control or 50% nuclear export compromised conditions, **c,** Representative still images of time-lapse movies of HeLa cells stably expressing H2B:GFP treated with either DMSO or 20 nM LMB at anaphase onset specifically. Scale bar, 10 μm. **d,** Chromatin-occupied area of H2B:GFP expressing HeLa cells treated with DMSO or 20 nM LMB at anaphase onset. Areas are normalized to nuclear cross-sectional area of DMSO treated cells at the anaphase to telophase transition (corresponding to ~ 10 minutes post anaphase onset) to compare nuclear expansion dynamics. Error bars represent SD.

**Figure 5.**
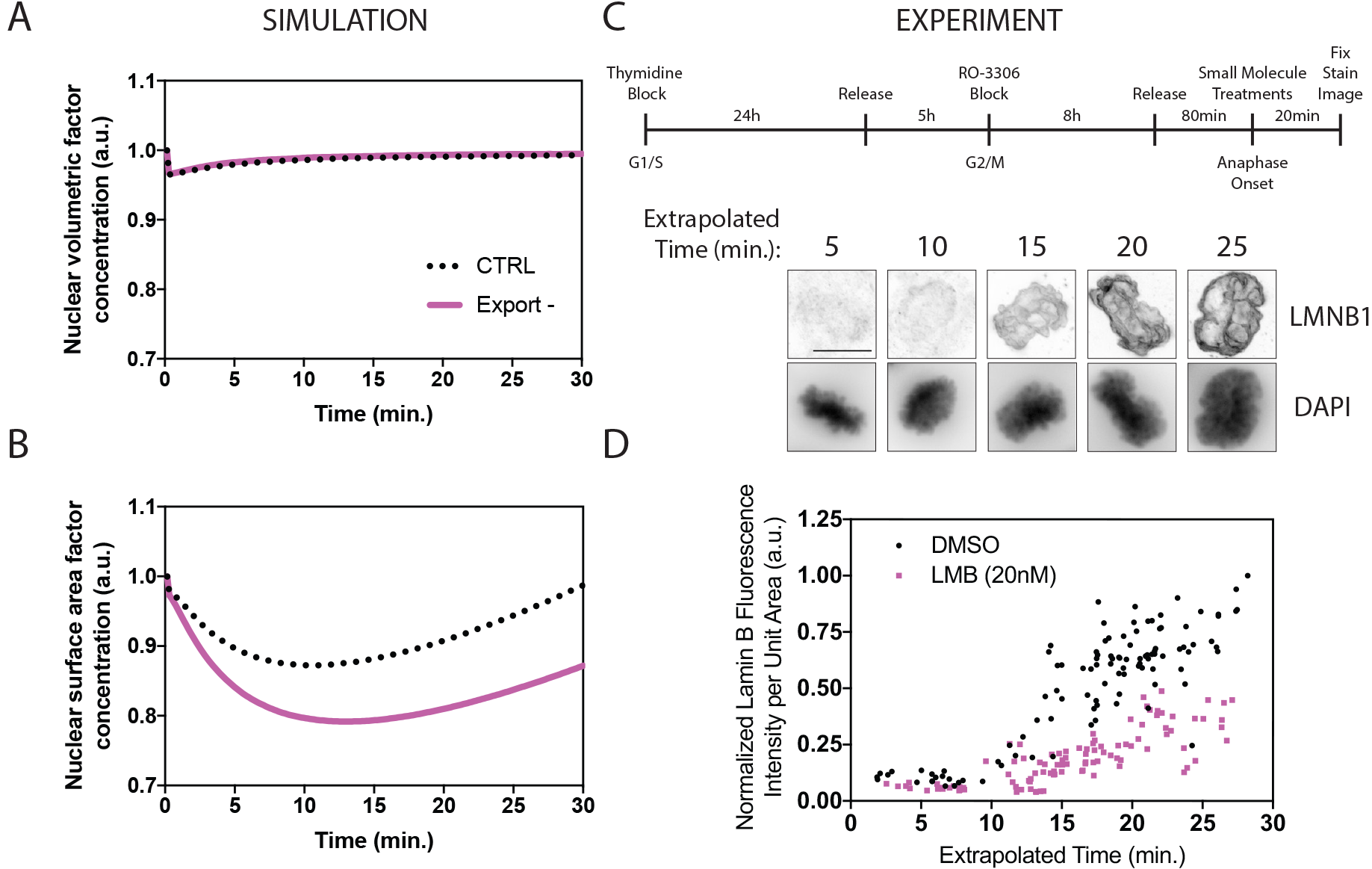
Inhibiting nuclear export during nuclear expansion reduces the nuclear concentration of LMNB1, a proposed nuclear surface area factor. **a,** Predicted nuclear concentrations of a proposed nuclear volumetric factor, NuMa. **b,** Predicted nuclear concentrations of a proposed nuclear surface area factor, LMNB1, through nuclear expansion in control and nuclear export compromised conditions. Nuclear export activity is compromised by 50% in the simulation. **c,** Representative still images of fixed HeLa cells synchronized in G2/M, released into mitosis, treated with small molecule inhibitors at approximately anaphase onset and fixed and stained for DAPI and LMNB1. Scale bar, 10 μm. **d,** Normalized LMNB1 fluorescence intensity in 3D maximum intensity projections proximal to DNA as highlighted by DAPI. Extrapolated times were determined for individual nuclei based on chromatin-occupied area (measured in Fig. 4D).

### Nuclear expansion is regulated by nucleo-cytoplasmic trafficking of nuclear volume and surface area factors

Given that nuclear surface area is not limited in multinucleate cytoplasmic extract droplets (i.e., Figure 1 does not agree with Figure 2), we set out to test how different levels of nuclear volumetric and surface area factors regulate the formation of the nucleus. We reasoned that once in proximity to the developing nucleus, these factors must be subject to NCT to enable their nuclear functions. Modifying NCT rates in the mathematical model detailed above and simulating nuclear expansion suggested that by inhibiting nuclear export by a factor of two the nucleus would expand further than its wild-type counterpart (Fig. 4A and 4B). The inverse effect is predicted by inhibiting nuclear import.

To test this prediction, HeLa cells stably expressing H2B:GFP were imaged as of anaphase-onset through mitotic exit – a transition in the cell cycle during which condensed mitotic chromatin must decondense to form an interphase-like nucleus. Cells were treated with inhibitors targeting nuclear import or export at anaphase onset specifically (Fig. 4C). A previously described area-based live-cell image analysis assay was adapted to measure the area occupied by chromatin as a measure of nuclear expansion (Maddox et al., 2006). To improve temporal resolution computationally, a temporal super-resolution method (Berro et al., 2014) was implemented. Inhibiting nuclear export with Leptomycin B (LMB) (Kudo et al., 1999) at anaphase onset (Fig. S1) induced further nuclear expansion as predicted (Fig. 4B), with nuclei reaching volumes approximately 50% larger than control nuclei (Fig. 4D). Inhibiting nuclear import with Importazole (Soderholm, et al., 2011) IPZ failed to display significant effects on nuclear expansion, which can be attributed to the compound’s subtle effects on nuclear import inhibition (Fig. S1). To exclude a role for NCT in regulating progression through mitotic exit globally, we developed an edge-based quantitative time-lapse microscopy assay to measure cell shape changes during cytokinesis in DIC images. Treating cells at anaphase with LMB did not significantly affect cell shape changes from metaphase through mitotic exit (Fig. S2).

In many animals, following fertilization of the oocyte the highly condensed sperm chromatin must expand to form an interphase-like pronucleus prior to pronuclear meeting (Matsumoto et al., 1999) - a process akin to nuclear expansion following mitotic exit in fully differentiated somatic cells. To test whether NCT regulates nuclear expansion of sperm chromatin following fertilization, we measured male pronuclear size following fertilization in the 1-cell C. elegans embryo. Compromising nuclear import or nuclear export by partially depleting IMB-1 (Importin β, 12 h treatment) and XPO-1 (Exportin-1, 24 h treatment) respectively, pronuclear expansion rates recapitulated our model’s prediction (Fig. S3).

Taken together, experimental inhibition of nuclear export during nuclear expansion yielded nuclei expanding further than their control counterparts as predicted by halving nuclear export rates of nuclear volumetric and surface area factors in silico.

### Surface area factor concentrations are regulated by nucleo-cytoplasmic trafficking

Although NCT regulates nuclear expansion rates, how nuclear volumetric and surface area factors contribute to this phenomenon was unclear. NuMa was modelled as a nuclear volumetric factor, as it is one of the most abundant proteins in the nucleus besides histones (Wühr et al., 2014). Our mathematical model predicted that the nuclear volumetric factor’s nuclear concentration was not severely perturbed by different NCT rates as the nucleus expanded (Fig. 5A). Conversely, when slowing nuclear import or export rates, the nuclear concentration of surface area factors was perturbed (Fig. 5B).

To test the prediction that a nuclear surface area factor must be trafficked between the cytoplasm and the nucleus during nuclear expansion, we measured the amount of endogenous Lamin B1 (LMNB1) during nuclear organogenesis using a fixed cell assay that allows temporal resolution using nuclear cross-sectional area as a predictor (Kafri et al., 2013). HeLa cells were synchronized at the G2/M transition using the CDK1 inhibitor RO-3306 (Vassilev et al., 2006) (Fig. 5C), released in mitosis and treated with small molecule inhibitors 80 minutes following their release into mitosis, approximately corresponding to anaphase onset, for 20 minutes. Cells were fixed and stained for Lamin B1 and DNA, which allowed the quantification of nuclear area occupied by chromatin. Using fitted curves describing the first 30 minutes of nuclear expansion following anaphase onset (Fig. 4D and Fig. S1E), timing of mitotic exit progression could be extrapolated computationally based on chromatin-occupied area of individual nuclei (Fig. 5C). Staining for endogenous LMNB1 revealed nuclear export inhibition limited the nuclear concentration of LMNB1 during nuclear expansion (Fig. 5D). Assuming LMNB1 is a core nuclear surface area component, reduced LMNB1 nuclear concentration following LMB treatment is consistent with our model’s prediction (Fig. 5B).

### Disrupting nucleo-cytoplasmic trafficking rates in Drosophila embryos affects nuclear function following nuclear organogenesis

Thus far, our data have suggested that balancing volumetric and surface area factors through NCT is important to build the nucleus. Despite the illustrated role for NCT in regulating nuclear organogenesis, it remains unclear how disrupting nuclear assembly affects downstream nuclear functions. Using a previously described approach to quantify transcription dynamics in the Drosophila early embryo (Garcia et al., 2013) (Fig. 6A), we tested how perturbing NCT in vivo affects nuclear function following nuclear assembly.

**Figure 6.**
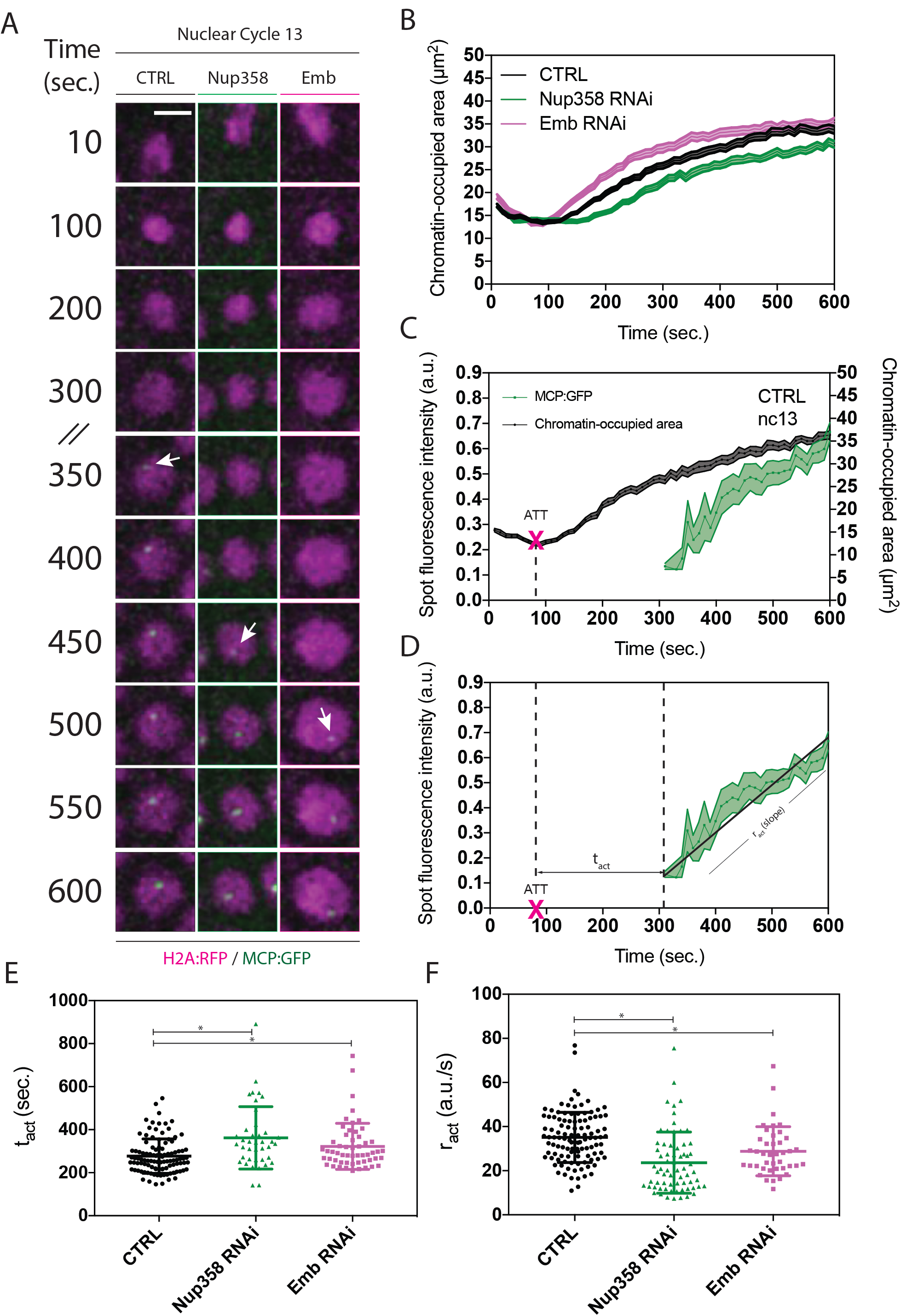
Depleting core nucleo-cytoplasmic trafficking components via RNAi affects the timing and rate of transcription activation in vivo. **a,** Representative still images of Drosophila embryos expressing H2A:RFP and MCP:GFP, which are being provided maternally, and a lacZ reporter gene controlled by the hb P2 enhancer and promoter, which is being provided paternally. Nup358 and Emb are targeted by a single copy of shRNA driven by a single copy of the mat-tub-Gal4 driver, both being maternally contributed. Images represent individual nuclei through embryonic nuclear cycle 13, where time 0 sec is anaphase onset. White arrows indicate MCP:GFP focus formation. Scale bar, 5 μm. **b,** Chromatin-occupied area of nuclei in control embryos, Nup358 depleted embryos and Emb targeted embryos. Areas are normalized to the expansion state of control embryos at the anaphase to telophase transition (corresponding to ~ 90 seconds post anaphase onset) to compare nuclear expansion dynamics. **c,** Chromatin-occupied area of nuclei in control embryos and normalized MCP:GFP fluorescence intensity of transcriptionally active spots in n>50 nuclei across n>3 embryos. Based on chromatin-occupied area dynamics, the anaphase-telophase transition (ATT) was computed in individual nuclei as the timing of nuclear expansion initiation. **d,** Transcription dynamics were measured by fitting linear regressions to MCP:GFP spot fluorescence intensities in individual nuclei. Transcription activation slopes were computed (r_act_) and the timing of transcription activation (t_act_) representing the time between when the linear regression attains a fluorescence intensity of zero minus the time of ATT. **e,** and **f,** tact and ract in control, Nup358 partially depleted and Emb partially depleted embryos. Error bars in b, c, d represent SEM and error bars in e and f represent SD. * = p < 0.05.

To monitor transcription, a lacZ reporter gene with MCP:G-FP-binding stem loops at its 5’ end was expressed paternally in the early embryo, driven by the activity of the hunchback P2 enhancer. The P2 enhancer is one of three enhancers responsible for establishing the endogenous hunchback expression gradient (Perry, et al., 2012) and was used in this study based on our extensive knowledge of this developmental expression pattern (Driever and Nüsslein-Volhard, 1989; Margolis, et al., 1995). Additionally, the morphogenetic protein Bicoid serves as the primary input for hb transcription regulation (He et al., 2008; Liu and Ma, 2011; Cheung et al., 2014). Bicoid-driven transcription can therefore be visualized as spots in nuclei in the anterior of the developing embryo (Fig. 6A). The probe’s fluorescence intensity, directly proportional to the amount of transcript being made, has been shown to be correlated to the rate of polymerase loading as nuclei enter interphase (Garcia et al., 2013).

Using the area-based assay described in Figure 3 to quantify nuclear compaction and expansion dynamics (Fig. 6B), transcription was measured with respect to mitotic events in individual nuclei during nuclear cycle 13 of the developing embryo (Fig. 6C and 6D). To compromise nuclear export and nuclear import, a transgenic RNAi approach was used to deplete the Crm1 fly homologue Embargoed (Emb) and Nucleoporin 358kD (Nup358), respectively. Transcription activation (tact) was measured as the time between the anaphase-telophase transition and the initiation of nuclear MCP:GFP focus formation, while transcription activation rates (ract) were measured as slopes of linear regressions of MCP:GFP recruitment to nuclear foci (Fig. 6D). The parameters tact and ract therefore represent the timing of transcription activation (initial polymerase loading) and the dynamics of transcription activation (rate of polymerase loading) respectively.

Depleting Emb and Nup358 in the early embryo using either one or two copies of the maternal alpha tubulin Gal4 (mat-tub-Gal4) driver resulted in titratable amounts of embryo hatch rates, suggesting these components can be targeted in the early embryo while allowing partial embryonic development (Fig. S4). Depleting the export component Emb resulted both in defective transcription activation dynamics in nuclear cycles 12 and 13 and in timing of transcription activation (tact) in nuclear cycle 13 (Fig. 6E). Similar effects on transcription activation dynamics and timing were observed in Nup358 depleted embryos. Phenotypes in transcription dynamics were more severe in Nup358 depleted embryos versus Emb depleted embryos (Fig. 6F), which is consistent with embryo hatch rates (Fig. S4). Interestingly, depleting Emb and Nup358 in embryos also recapitulates our mathematical model’s predicted effect of nuclear import/ export dynamics on nuclear expansion. By normalizing nuclear expansion values to maximally condensed chromatin at the anaphase-telophase transition in control embryos, nuclear expansion dynamics could be compared between experimental conditions as nuclei reassemble during development. Inhibiting nuclear import by partially depleting Nup358 induced slower nuclear expansion whereas inhibiting nuclear export by partially depleting Emb induced further nuclear expansion rates during nuclear cycle 13 of developing Drosophila embryos (Fig. 6B), further validating the predicted role for nuclear import/export-regulated nuclear expansion.

These results suggest that disrupting either nuclear import or nuclear export, which we have shown disrupts nuclear assembly, affects downstream nuclear functions such as transcription in a developing organism.

## DISCUSSION

Combining *in vitro*, *in silico* and *in vivo* predictions and observations across several cell biological contexts, we uncover novel regulatory mechanisms essential to nuclear organogenesis and to downstream nuclear function. In frog egg extracts, explicitly dissecting nuclear volume, nuclear surface area and cell size scaling relationships by modulating the number of nuclei in a defined cytoplasmic volume, we determined that neither nuclear surface area nor nuclear volume scale directly with cell size (Fig. 2). Seeking to explain this phenomenon and how nuclei expand to given sizes, we developed a mechano-chemical mathematical model (Fig. 3) based on prior biochemical knowledge (Wühr et al., 2014) that is dependent on microtubule and dynein-based cargo transport (Fig. 2C, D), nucleo-cytoplasmic trafficking, hydrostatic pressure, osmotic pressure and nuclear envelope surface tension. Our model recapitulates nuclear expansion (Fig. 3C) and the scaling relationship between nuclear surface area and cell size *in vitro* (Fig. 3D).

In silico, our model predicted NCT was a key nuclear expansion regulatory mechanism (Fig. 4A and 4B). Treating HeLa cells at anaphase onset specifically with the nuclear export inhibitor LMB and using a previously described area-based image analysis approach, we uncover a novel role for nuclear export in regulating nuclear expansion (Fig. 4C and 4D). Using IPZ to target nuclear import subtly affected the import of an NLS:mCherry reporter (Fig. S1C), but did not recapitulate our model’s prediction in HeLa cells (data not shown). We postulate that our inhibition of nuclear import using IPZ was partial and insufficient to generate a nuclear expansion phenotype. This hypothesis is consistent with RNAi-based perturbations of Impβ, which have also been shown to partially affect Golgi clustering in anaphase in mammalian cells (Schmitz et al., 2010). Our model also predicted that NCT modulated the nuclear accumulation of a nuclear surface area factor as opposed to a nuclear volumetric factor (Fig. 5A and 5B). By staining for endogenous LMNB1 (a proposed nuclear surface area factor) and using a computational method to extrapolate time based on area occupied by chromatin (Kafri et al., 2013), we determine that inhibiting nuclear export reduces the amount of nuclear LMNB1 during nuclear expansion (Fig. 5C and 5D). Given the counter-intuitive nature of LMNB1 regulation by nuclear export, we hypothesize that negative regulators must be timely exported from the reforming nucleus to allow proper LMNB1 localization. Such biochemical factors could be negative PP2A-B55 regulators, which is known to regulate anaphase Golgi clustering (Schmitz et al., 2010), nuclear pore assembly and nuclear import (Cundell et al., 2016). Identifying the factors that must be exported from the reforming nucleus will be crucial in understanding the molecular requirements for proper nuclear organogenesis.

To determine whether disrupting NCT affected nuclear function following organogenesis, we measured transcription activation timing and rates with respect to nuclear expansion in individual nuclei in developing fly embryos (Fig. 6). First, partially depleting Nup358 (targeting import) slowed nuclear expansion, while depleting Emb (targeting export) accelerated expansion during nuclear cycle 13 (Fig. 6D), as predicted by our model (Fig. 4B). Second, targeting either nuclear import or nuclear export delayed timing of transcription activation in addition to slowing transcription activation dynamics (Fig. 6E). Although transcription activation dynamics were used as a readout for nuclear function in this study, it is likely that numerous nuclear functions including chromatin organization and re-establishing epigenetic modifications would be affected in the interphase following an abrogated nuclear assembly. Taken together, our in vivo observations support a role for NCT in regulating nuclear expansion, a role that is consequential in downstream nuclear function.

Despite our understanding of many of the spatial, temporal and regulatory aspects of nuclear envelope components such as LMNB1, trafficking components such as Impβ and mitotic kinases such as Aurora B and CDK1, our understanding of how these components function as whole to regulate nuclear expansion and nuclear organogenesis has remained unclear. Our model and the uncovered role for NCT in regulating nuclear expansion and downstream nuclear function by limiting nuclear surface area factors provides novel insight into how the nucleus is built. Evidently, building a functional nucleus must require several components and timely regulatory mechanisms that extend beyond the scope of this study. As this simple mechano-chemical model was used to predict NCT’s role in regulating nuclear expansion, this model could be used to address a variety of important questions including how microtubule-based transport of different factors regulates nuclear expansion. In fact, given the emerging evidence that microtubule targeting agents (MTAs) used clinically as chemotherapeutic agents largely fail to target mitotic cells in vivo (discussed in Komlodi-Pasztor et al., 2011), this model may prove useful in predicting and testing a role for MTAs in regulating interphase nuclear metabolism.

In expanding on our model and its insight into nuclear expansion, several other mechanisms could be included and implemented to improve its accuracy in predicting biological phenomena and generating hypotheses. For example, once nuclear surface area factors reach the nucleus and are imported, mechanisms regulating their assembly at the nuclear envelope may play equally important temporal roles in nuclear assembly. Incorporating regulatory mechanisms such as lamina assembly is likely to improve hypotheses generated from the model and provide insight into how nuclei come to be.

## METHODS

### Xenopus extract use, microfluidics and microscopy

Cytostatic factor (CSF)-arrested egg extracts were prepared as described previously (Desai et al. 1999; Hannak and Heald, 2006). Interphase extract was maintained by the addition of calcium and cyclohexim-ide. De-membranated sperm nuclei and fluorophores were subsequently added to the extract, which was immediately loaded into microfluidic devices. m-Cherry HURP or GFP-NLS were added to the extract at 2 uM to visualize nuclear import. The standard nuclear assembly reaction was 100 μl fresh extract, 0.4 mM CaCl2, 100 μg/ml cycloheximide and 1000 Xenopus sperm per μl. Reactions were incubated at 16-18°C and spherical, import-competent nuclei generally formed within 30-45 min.

Microfluidic devices were cast in polydimethylsiloxane (PDMS, Sylgard 184, Dow Corning) using well-established soft lithography techniques, which are described in detail elsewhere (Duffy et al., 1998; Xia & Whitesides, 1998). Briefly, a microchannel network was designed in AutoCAD (Autodesk, Inc) drafting software. The photomask was output to film (CAD/Art Services, Bandon, OR) and used to lithographically expose the network in a photoresist film (SU-3025, MicroChem, Newton, MA) spun onto a silicon wafer at 30 μm. Polydimethylsiloxane (PDMS) elastomer was then poured over the patterned wafer, cured at 70°C, and removed, producing an imprinted channel network. Fluid inlet and outlet ports were punched into the PDMS with a sharpened, unbeveled syringe tip (Brico Medical, Dayton, NJ). The PDMS channel network was exposed to oxygen plasma (Harrick Plasma, Ithaca, NY) and placed in contact with a cover glass slip (#1.5, Thomas Scientific) to form an irreversible bond.

### Mathematical model

The mathematical model is derived from a combination of mechanical force-balance determining the size of the nucleus, and transport of surface and volume factors throughout the cytoplasm and into the nucleus. The almost-spherical shape of in vitro nuclei allows us to impose spherical symmetry on the nucleus and describe its size with a single variable, the radius r(t). Mechanical force-balance thus becomes a single ordinary differential equation (ODE) for dr/dt. To describe transport in the cytoplasm, we make the simplifying assumption that it can be divided into two compartments, “proximal” to the nucleus and “distal” to the nucleus. This leads to 6 ODEs: for proximal, distal and nuclear amounts of surface and volume factors. The full model is thus a coupled system of 7 nonlinear ODEs. These equations are solved numerically in Matlab (Mathworks). There are 15 biophysical parameters in the model. Of these, 9 are taken from independent experiments or estimated from previous work. The remaining 6 (nuclear import and export rates and initial amounts of both factors) are fit to the time series for growth, as described in the Supplemental Math.

### Mammalian tissue culture cell maintenance, drug treatments, immunostaining and microscopy

HeLa cells stably expressing H2B:GFP were cultured in DMEM (containing 1g/L glucose) supplemented with 10% fetal bovine serum and penicillin-streptomycin (Sigma) at 37°C in a humidified 95% air, 5% CO_2_ incubator. To target nuclear import, the small molecule inhibitor Importazole (IPZ) that targets the interaction between Ran-GTP and Importin-ß was used at concentrations consistent with previously published results (Soderholm et al., 2011). To target nuclear export, Leptomycin-B (LMB), an irreversible inhibitor of the main nuclear protein export component Crm1, was used (Kudo et al., 1999). Immunostaining was performed using a 5 minute 0.2% TritonX-100 permeabilization, followed by a 20-minute fixation with 4% EM-grade paraformaldehyde (Ted Pella) in PHEM. All steps are performed with coverslips in tissue culture dishes placed on top of a hot plate heated to 37°C. LMNB1 was stained with a rabbit polyclonal antibody from Proteintech (#12987-1-AP).

Live-cell microscopy of tissue-culture cells was performed at 37°C in a humidified chamber on a DeltaVision microscope using Softworx software (Applied Precision) with a CoolSnap HQ2 camera (Photometrics) and a x60 planApo objective. Cells were incubated in CO_2_-independent media (ThermoFisher) in 6-channel μ-Slides (Ibidi) during image acquisition. Fixed-cell microscopy was performed on the same DeltaVision microscope as mentioned above at room temperature.

To measure nuclear import or nuclear export, an NLS or an NES respectively was cloned into pOD69 with an mCherry tag as to obtain fusion proteins containing either the NLS or the NES with two fluorophores to prevent diffusion across nuclear pore complexes. The SV40 NLS (PK-KKRKV) was used to follow Impβ-directed nuclear import (Kalderon et al., 1984). HIV-1’s viral Rev protein’s NES (LPPLERLTL) was used to follow nuclear export (Fischer et al., 1995; Neville et al., 1997).

### *C. elegans* use, RNAi and microscopy

The worm strain TH32 (pie-1∷bg-1∷GFP + unc-119(+), pie-1∷G-FP∷H2B + unc-119(+)), was grown and maintained at 20°C using standard procedures. Bacterial strains containing a vector expressing dsRNA under the IPTG promoter were obtained from the Ahringer library (from Bob Goldstein’s laboratory, University of North Carolina at Chapel Hill, Chapel Hill, NC, USA). Targets were confirmed by sequencing.

Worm embryos were mounted in Egg buffer (118 mmol/L NaCl, 48 mmol/L KCl, 2 mmol/L CaCl_2_, 2 mmol/L MgCl_2_, 25 mmol/L HEPES, pH 7.3) between a 1.5 coverslip and a microscope slide spaced by 22.81μm glass beads (Whitehouse Scientific) and sealed with Valap (1:1:1 lanolin, petroleum jelly, and parafilm wax). Embryos were then imaged on a Nikon A1r resonant scanning confocal microscope using a x60 Apo water immersion objective (Nikon), GaASP PMT detectors and NIS-Elements (Nikon) at 22°C.

### Drosophila husbandry and microscopy

Female virgins maternally expressing MCP:GFP either UAS-emb RNAi (TRiP #HMS00991) or UAS-nup358 RNAi (TRiP #HMS00865) with males expressing H2A:RFP and mat-tub-Gal4. Female virgins expressing single copies of MCP:GFP, H2A:RFP, mat-tub-Gal4 and either UAS-emb or UAS-nup358 were obtained and subsequently crossed with males of the reporter line (hb P2 enhancer and promoter). Flies expressing single copies of MCP:GFP and H2A:RFP were used as controls. Embryos were dechorionated, mounted between gas permeable lumox^®^ dish 50 dishes and a coverslip and imaged on a Nikon A1r resonant scanning confocal microscope. Images were acquired using a x60 Apo TIRF objective. All fly husbandry, crosses and imaging were performed at 22°C.

### Image analysis

Nuclear expansion, transcription activation and DIC morphology assay ImageJ plugins are available on VB’s GitHub page (https://github.com/viboud12). Other image analysis plugins used in nuclear import/export, pronuclear size and droplet nuclear size analysis are available upon request.

## SUPPLEMENTAL INFORMATION

Supplemental Information includes Supplemental Math and four figures and can be found with this article.

## ACKNOWLEDGMENTS

We greatly thank Chris Obara and Abby Buchwalter for providing reagents. We also thank Wallace Marshall for and the support of the NSF in organizing the Cell Modeling Hackathon in Half Moon Bay, CA. during which this collaborative effort was initiated. Furthermore, we would like to thank Vincent Archambault and Talia Hatkevich for critical reading of the manuscript. V.B. was supported by predoctoral fellowships from the Fonds de Recherche Santé-Québec (FRQS). This study was also supported by National Science Foundation CAREER Award 1652512 to P.M., NIGMS grant R01GM113028 to J.G. and NSF DMS 1454739 to J.A. Transcription activation experiments were supported by post-course research funds provided to V.B. through the Marine Biological Laboratory’s Physiology course.

## CONTRIBUTIONS

V.B. designed and carried out experiments in mammalian tissue culture, *C. elegans*embryos and Drosophila embryos, wrote ImageJ-based image analysis code, performed image analysis and wrote the manuscript. J.H. and P.C. carried out Xenopus egg extract experiments and J.H. made initial nuclear surface area to volume observations. J.S. implemented the temporal super-resolution method inspired by (Berro and Pollard, 2014). R.R. Wrote code for simulations, ran simulations to explore model variants and parameters, and generated data for figures. K.M. Developed models, helped write code for simulations, explored model variants and generated data for figures. H.G. contributed to transcription activation experimental design and analysis. V.B. wrote the manuscript with significant contributions from J.A.

## Mathematical model of nuclear assembly

### 1 Basic model equations and assumptions

#### 1.1 Mechanics of the nucleus

The nuclear volume *V*(*t*) and surface area *S*(*t*) are both dynamic quantities. The total mechanical free energy of the nucleus, which determines nuclear shape dynamics, is itself determined by several factors. As our starting point, we assume that this energy has three dominant terms:

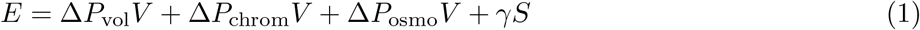

where *P*_vol_ is the hydrostatic pressure due to the hypothesized elastic volumetric network, *P*_chrom_ is the pressure due to chromatin, *P*_osmo_ is osmotic pressure, and *γ* is the surface tension, which includes the full nuclear envelope (membranes and lamina). These are shown in Fig. S1 (Bottom). This energy drives changes to nuclear volume according to

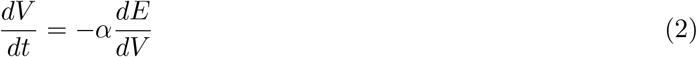

where *α* is the hydraulic permeability [S5] of the nucleus, which has units of nm^3^/(Pa · s). In words, Eq. 2 states that the nuclear volume will increase or decrease in order to minimize the free energy, at which point the three forces in Eq. 1 are in balance.

**Fig. S1.**
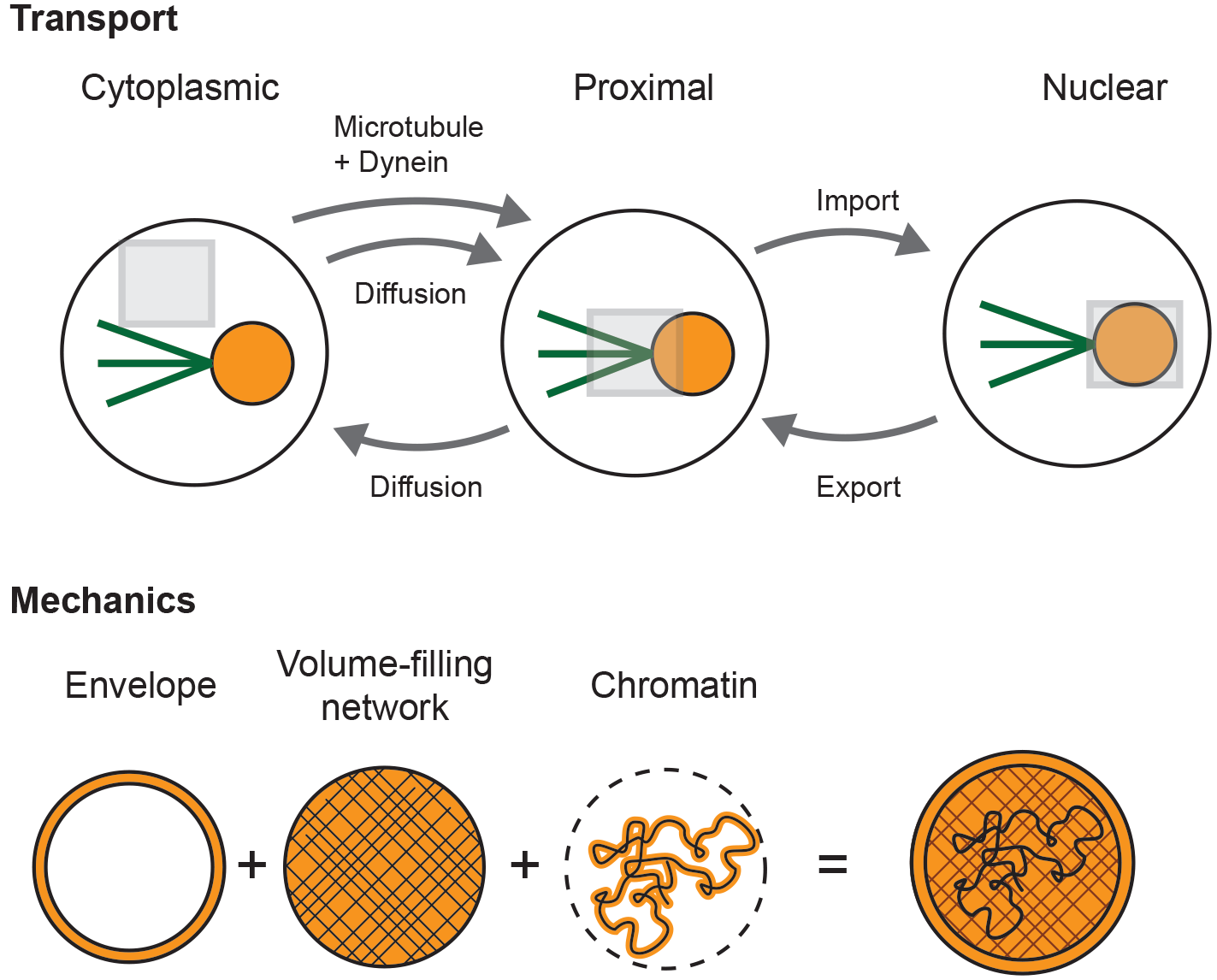
(Top) Transport of nuclear assembly factors. Simplified model with three region. Same transport mechanism is assumed for both the volume factor and surface factor. (Bottom) Mechanical factors included in model that (a priori) can determine nuclear size.

In general, the nucleus might buckle into a non-spherical shape. However, following data here, we first formulate the model assuming it remains approximately spherical. This approximation is valid provided we are below the buckling threshold [S5]. Combining Eq. 1, 2, and the spherical approximation, we obtain

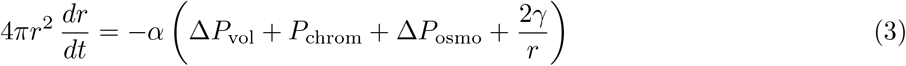

where *r*(*t*) is the nuclear radius. In absence of precise measurements of the permeability *α*, we make the reasonable assumption that *α* is proportional to the surface area, *α* = *α*_0_4π*r*^2^ (canceling the left-hand-side prefactor).

Pressure due to confining chromatin has been estimated to satisfy [S6, S8]

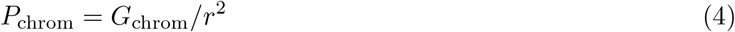

where *G*_chrom_ is the chromatin compressibility modulus. We treat both the bulk inside the nucleus and the nuclear envelope as elastic materials. Therefore, they have elastic compressibility *G*_vol_ and extensibility *G*_surf_, and the pressure and surface tension are given by

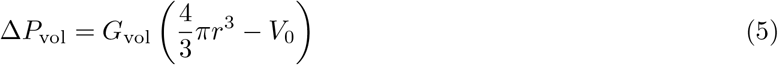

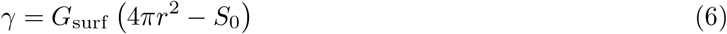

where *V*_0_(*t*) and *S*_0_(*t*) are the preferred volume and surface area, i.e., the “rest” volume and “rest” area. Note that, *a priori*, we do not assume which term dominates, i.e., it could be that *G*_surf_ is negligible compared to the bulk volume compressibility.

#### 1.2 Transport of nuclear assembly factors

During nuclear assembly, material is added to the nucleus via regulated import at nuclear pore complexes (NPCs). This includes nuclear material destined to assemble throughout the nucleoplasm, such as NuMA [S7], and material destined to assemble the envelope, such as lamin-A/C and lamin-B [S2]. We generically refer to these as a nuclear volume factor *f*^vo1^ and nuclear surface factor *f*^surf^.

At the beginning of nuclear assembly, these factors have a distribution throughout the cytoplasm. As a first approximation, we divide the cytoplasmic space into two regions: distal to the nucleus and proximal to the nucleus, with volumes *V*_cyto_ and *V*_prox_ respectively. This is shown schematically in Fig. S1 (Top).

Material is transported between these regions by two mechanisms: passive diffusion, which is bi-directional, and active transport by dynein along *N*_*MT*_ centrosomal microtubules with velocity *υ*_dyn_. Once the factor is proximal to the nucleus, its exchange with the nucleus is regulated by NPCs with first-order kinetic rates *k*_in_ and *k*_out_ (both with units of *μ*M^−1^ s^−1^ per NPC). This leads to dynamic equations

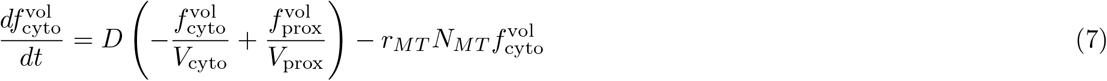

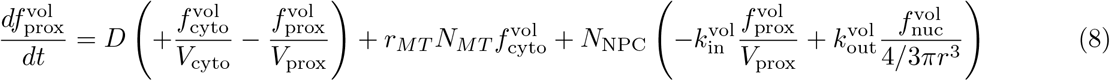

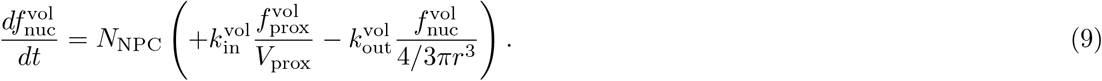

The diffusion parameter 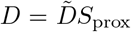 is the molecular diffusion coefficient (with units of nm^2^/ s) times the area of interface between the proximal cytoplasmic region and distal cytoplasmic region. The active transport parameter *r*_*MT*_ = *υ*_dyn_/*l*_prox_ is the dynein velocity divided by the mean transport distance between distal and proximal cytoplasmic regions. These definitions of *D* and *r*_*MT*_ assume each molecule of factor is equally likely to be transported actively or by diffusion. We can adjust this assumption by decreasing either (or both) of these parameters.

Similarly for surface factor,

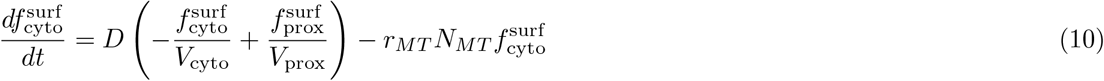

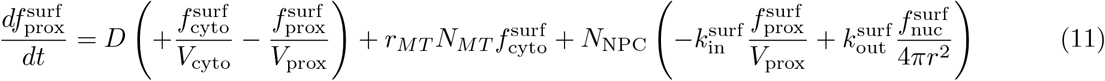

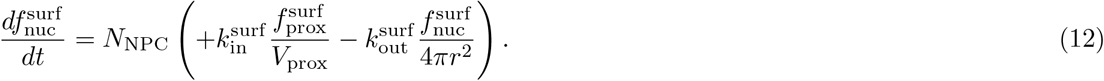

Note that the nuclear amount of factor 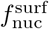 scales with nuclear surface area, rather than nuclear volume.

### 2 Parameter estimation and model simplification

Instead of tracking factor amounts in numbers of molecules, we define 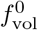 as the amount (i.e., number of molecules) of volume factor required to assemble one unit of nuclear volume, and 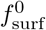 as the amount of surface factor required to assemble one unit of nuclear surface, and set 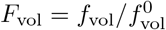 and 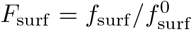. This allows us to re-write the force-balance equation Eq. 3 as

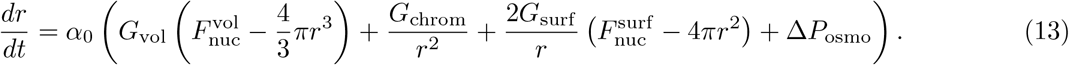

The units of *k*_in_ and *k*_out_ are now s^−1^.

#### 2.1 Diffusional delivery rate in cytoplasm to nuclear periphery

We split the cytoplasm into two regions, which we term the cytoplasm (distal to the nucleus) and the proximal region (the nuclear periphery). Diffusive transport between two regions obeys the diffusion equation that simplifies, for the case of two compartments, to

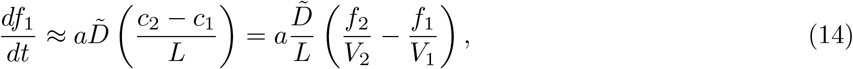

where 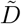 is the true diffusion coefficient, estimated to be 10*μ*m^2^/*s* [S4], and *c*_*i*_ is the concentration, *f*_*i*_ is the factor, and *V*_*i*_ is the volume of a given region. *L* denotes the length over which factor is transported and *a* is the cross-sectional area separating the regions. To apply this to our cytoplasmic systems, we define L ≈ *r*_ce11_ and 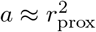. This leads to

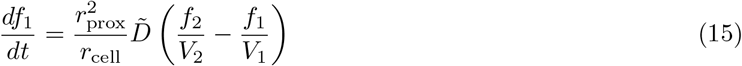

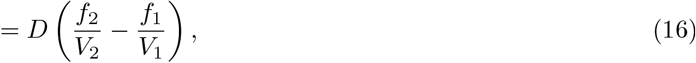

where we have defined the diffusion rate parameter

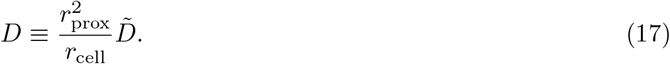

**Table S1.** Model parameters.

| Parameter | Description | Values for Xenopus | ... for HeLa | Source / note |
| --- | --- | --- | --- | --- |
| $\alpha_0$ | Per-area hydraulic permeability | $10^{-6} \mu\text{m}/\text{Pa} \cdot \text{min}$ | $10^{-6} \mu\text{m}/\text{Pa} \cdot \text{min}$ | [S5], see Sec. 2.2 |
| $D$ | Diffusion rate parameter | $10^3 \text{s}^{-1}$ | $10^3 \text{s}^{-1}$ | Estimated in Sec. 2.1 |
| $N_{MT}$ | Number of microtubules involved in transport | 50 | 50 | Observed here |
| $r_{MT}$ | Transport rate for microtubule-based delivery | $10^{-3} \text{s}^{-1}$ | $3 \times 10^{-4} \text{s}^{-1}$ | Fit in Sec. 3.3 |
| $G_{\text{vol}}$ | Volumetric factor bulk modulus | $10^4 \text{Pa}/\mu\text{m}^3$ | $5 \times 10^6 \text{Pa}/\mu\text{m}^3$ | Fit in Sec. 3.2 |
| $G_{\text{chrom}}$ | Chromatin confinement coefficient | $10^9 \text{Pa} \mu\text{m}^2$ | $10^9 \text{Pa} \mu\text{m}^2$ | Fit in Sec. 3.2 |
| $G_{\text{surf}}$ | Surface factor elastic modulus | $5 \times 10^3 \text{pN}/\mu\text{m}^3$ | $5 \times 10^3 \text{pN}/\mu\text{m}^3$ | Fit in Sec. 3.2 |
| $V_{\text{cyto}}$ | Volume of cytoplasm, distal region | $6.5 \times 10^4 (\mu\text{m}^3)$ | $4 \times 10^3 \mu\text{m}^3$ | Measured |
| $V_{\text{prox}}$ | Volume of nuclear proximal region | $3 \times 10^3 \mu\text{m}^3$ | $10^3 \mu\text{m}^3$ | Arbitrary |
| $N_{NPC}$ | Number of nuclear pore complexes | $3 \times 10^3$ | $3 \times 10^3$ | [S1] |
| $k_{\text{in}}^{\text{surf}}$ | Import rate of surface factor | $0.25 \text{s}^{-1}$ | $10^{-3} \text{s}^{-1}$ | Fit in 3.4 |
| $k_{\text{out}}^{\text{surf}}$ | Export rate of surface factor | $0.05 \text{s}^{-1}$ | $10^{-6} \text{s}^{-1}$ | Fit in 3.4 |
| $k_{\text{in}}^{\text{vol}}$ | Import rate of volume factor | $0.65 \text{s}^{-1}$ | $8 \times 10^{-2} \text{s}^{-1}$ | Fit in 3.2 |
| $k_{\text{out}}^{\text{vol}}$ | Export rate of volume factor | $0.05 \text{s}^{-1}$ | $8 \times 10^{-3} \text{s}^{-1}$ | Fit in 3.2 |
| $F_{\text{tot}}^{\text{vol}}$ | Total amount of volume factor | $8 \times 10^3 \mu\text{m}^3$ | $1.5 \times 10^3 \mu\text{m}^3$ | Estimated |
| $F_{\text{tot}}^{\text{surf}}$ | Total amount of surface factor | $1.1 \times 10^4 \mu\text{m}^2$ | $1.1 \times 10^4 \mu\text{m}^2$ | Estimated |

Using *r*_prox_ ≈ 10−100*μ*m and r_ce11_ ≈ 100−10^3^*μ*m, we obtain *D* = 10^4^*μ*m^3^/min. This leads us to estimate that *D* = 10^3^ − 10^5^*μ*m^3^/min.

#### 2.2 Hydraulic permeability of nuclear envelope

To estimate the per-area hydraulic permeability *α*_0_, we refer to Kim et al. [S5], where *α* is a constant of proportionality between change in volume and pressure, i.e., *dV*/*dt* = *α*Δ*P*. They find parameter values *V*_*N*0_ = 805*μ*m^3^, 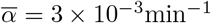, and *μ* = 10^4^ Pa, related to the permeability by their equation

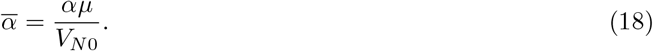

If the nucleus is spherical, as we assume in this model, the per-area hydraulic permeability is *α* = *α*_0_4π*r*^2^. Combining this with equations

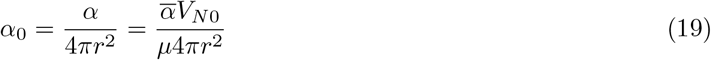

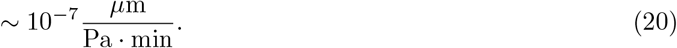

We find that this value of *α*_0_ is large enough that water import and mechanical equilibration is fast enough (during assembly) that nuclear growth is limited by other factors, specifically transport of material to the nuclear periphery and import of this material into the nucleus.

### 3 Simulation results

#### 3.1 Numerical simulation of model and fitting to experimental data

We simulate the model, which is a systems of 7 ordinary differential equations given by Eq 13 and 7-12, with 15 parameters. We begin simulating at *t* = 0, which we identify as the anaphase-to-telophase transition, after the nuclear envelope is sufficiently formed so that transport in and out is regulated via nuclear pore complexes. At this time, we observe that the nucleus has size *r*(0) = *r*_0_ = 4.18 *μ*m. We assume that it is in mechanical equilibrium with 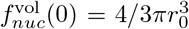 and 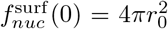 and that an equal concentration of the remaining factor are in the proximal and distal cytoplasmic pools. Of the 15 parameters, 6 are well-constrained by direct observation or by estimated from the literature, see Table S1. We explore ranges for the remaining 8 parameters as discussed in the sections below. From the seven dynamic variables (nuclear radius, and surface factor and volume factor amounts in each of three spatial compartments), we can further compute nuclear surface area, cross-sectional area and volume, and the concentration (amount per volume) of surface and volume factors. A sample time series produced by the simulation is shown in Fig. S2.

**Fig. S2:**
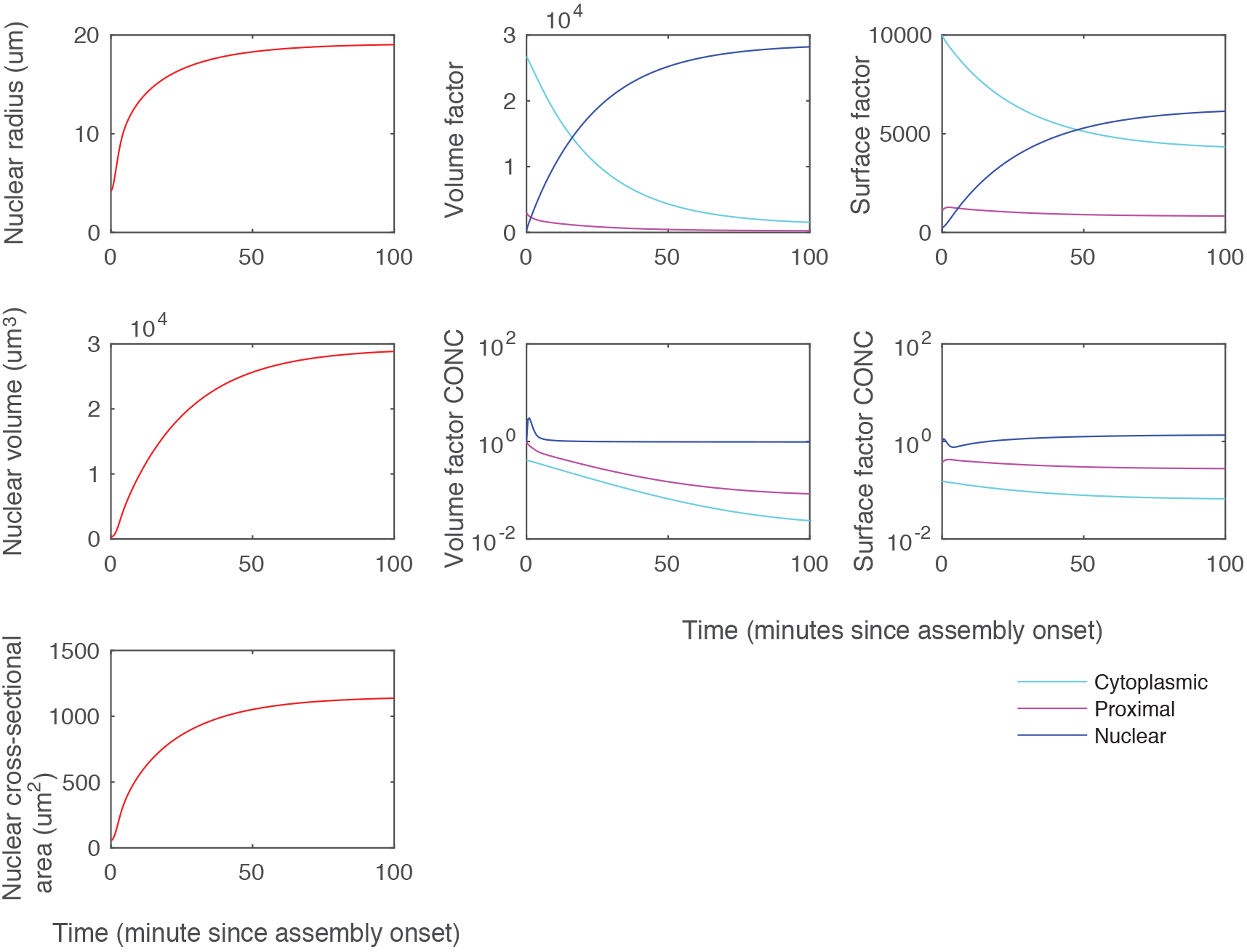
Sample time series produced by the simulation.

#### 3.2 Continuum of behavior between volume-dominated and surface-dominated regimes

We modify the model so that *N*_nuc_ nuclei share a common peripheral space and common pools of surface and volume factors. We then simulate the model until it reaches a steady state. The parameters with the least-well constrained values are the three mechanical moduli, specifically the surface factor compressibility *G*_vo1_, the volume factor compressibility *G*_surf_, and the chromatic confinement coefficient *G*_chrom_, so we perform exploration in these three parameters.

We first explore the model under Hypothesis 2, i.e., that nuclear size is determined by a competition between volume and surface factors. We do this by setting the chromatin confinement coefficient *G*_chrom_ = 0. Resulting nuclear scaling is shown in Fig. S3. When the volume factor modulus is low compared to surface factor modulus (left top and bottom), the total nuclear surface area is constant. This necessarily implies a decreasing total nuclear volume. When the volume factor modulus is high compared to the surface factor modulus (right top and bottom), the total surface volume is constant. This necessarily implies an increasing total nuclear surface area. Note that this model cannot give rise to total nuclear volume that increases with number of nuclei. For this reason, our simulations argue against Hypothesis 2.

**Fig. S3:**
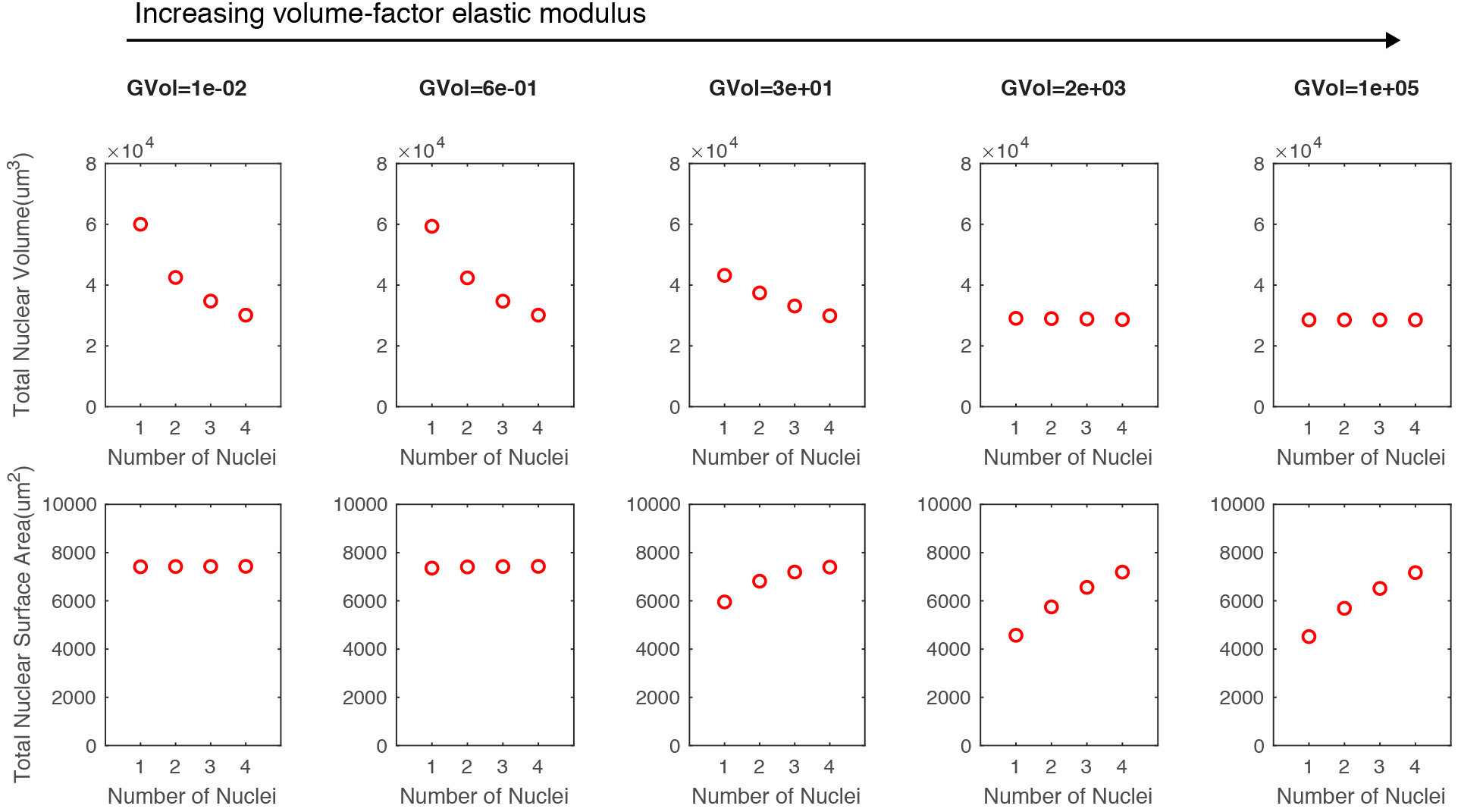
Nuclear size scaling as the ratio of *G*_vol_ (units of Pa/ *μ*m^3^)) to *G*_surf_ (pN/ nm^3^), with no chromatin effect (*G*_chrom_ = 0).

We next simulate the model under Hypothesis 3, i.e., a competition between surface factor, volume factor and chromatin, as shown in Fig. S4. To match the nuclear sizes in Main Text Figure 2, we find best-fit parameters shown in Table 1. In agreement with the finding of the Hypothesis 2 model, we find that agreement arises when the surface factor modulus is low, and therefore nuclear size is primarily determined by the remaining two factors, volume factor and chromatin.

To match the growth speed, i.e., the approach to this steady state, observed in Xenopus, we find that
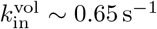. Note that the growth speed is approximately independent of 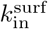 since the system is close to the volume-limited regime.

**Fig. S4:**
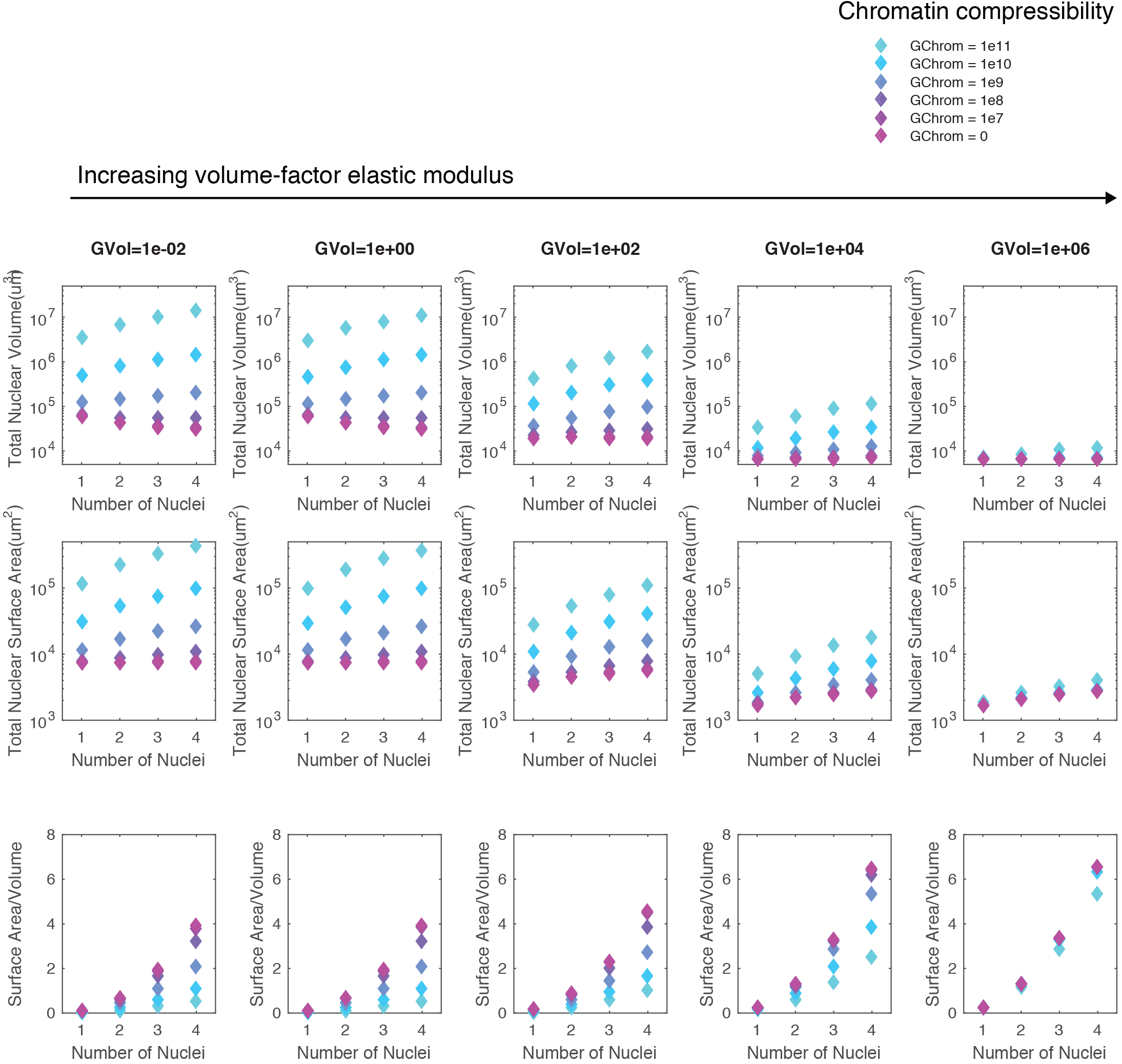
Nuclear size scaling with *G*_chrom_ = 0 (magenta) and for increasing *G*_chrom_ (units of Pa · *μ*m^2^, color axis). Note the log scale.

#### 3.3 Roles of diffusive and active delivery to nuclear periphery

Experiments have shown that inhibition of microtubule/dynein-based transport, e.g., using nocodazole, reduce the nuclear volume by approximately half [S3]. Roughly speaking, this suggests that half of delivery to the nuclear periphery is based on this active mechanism and, in the context of this model, suggests the remaining half of delivery is due to passive diffusion. In Fig. S5, we simulate the model with *N*_MT_ = 0 to mimic the nocodazole experiment (red curve and bar). We then simulate the control *N*_MT_ = 50 and find that, to reproduce the two-fold change, we must assume a per-microtubule delivery rate of *r*_*MT*_ 10^−3^ s^−1^.

**Fig. S5.**
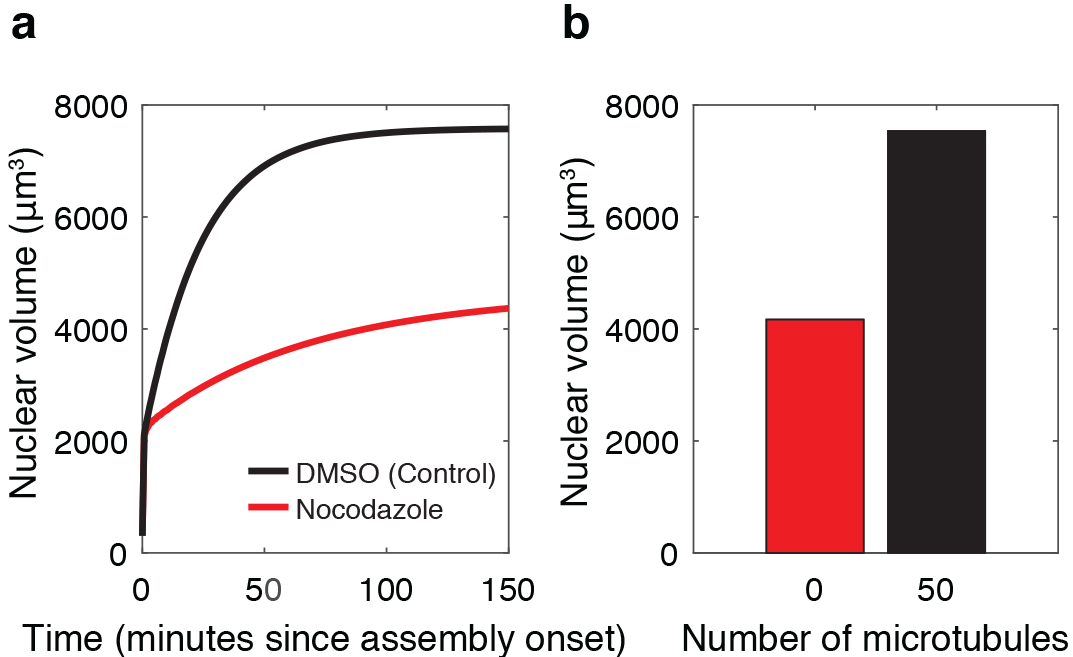
Microtubule/dynein-based transport inhibition imply relative importance of diffusive to active transport of material to nuclear-proximal region. (a) Steady-state nuclear size with and without microtubule/dynein-based transport. This constrains the parameter for microtubule/dynein transport rate, *r*_*MT*_ 10^−3^ s^−1^. (b) Time series of nuclear volume with and without microtubule/dynein-based transport.

#### 3.4 Nuclear export inhibition in HeLa

Finally, we simulate the inhibition of nuclear export through nuclear pore complexes by reducing 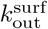 and 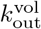 by half. In agreement with intuition, this leads to larger nuclear size, as shown in Main Text Fig. 4. However, counter-intuitively, this leads to a decrease in the concentration of surface factor in the nucleus, as shown in Main Text Fig. 5.

We note that although the time series shown are for specific choices of parameters, this behavior is generic under two conditions: First, if the nucleus is volume-limited, and second, if the the surface factor is import-dominated or, in the language of biochemistry, has a high affinity for the nucleus. In these two cases, reducing export leads to a larger nucleus because more volume factor is imported. Meanwhile, since all of the surface factor was already in the nucleus, the export inhibition cannot lead to more nsurface factor. Instead, the same amount is distributed over a larger volume, thus reducing its concentration.

At these parameters, at early times, the concentration of surface factor initially drops. At these times, the nucleus is small, therefore relatively small increases in size lead to dilution of surface factor and thus decrease in its concentration.

---

**Figure S1.**
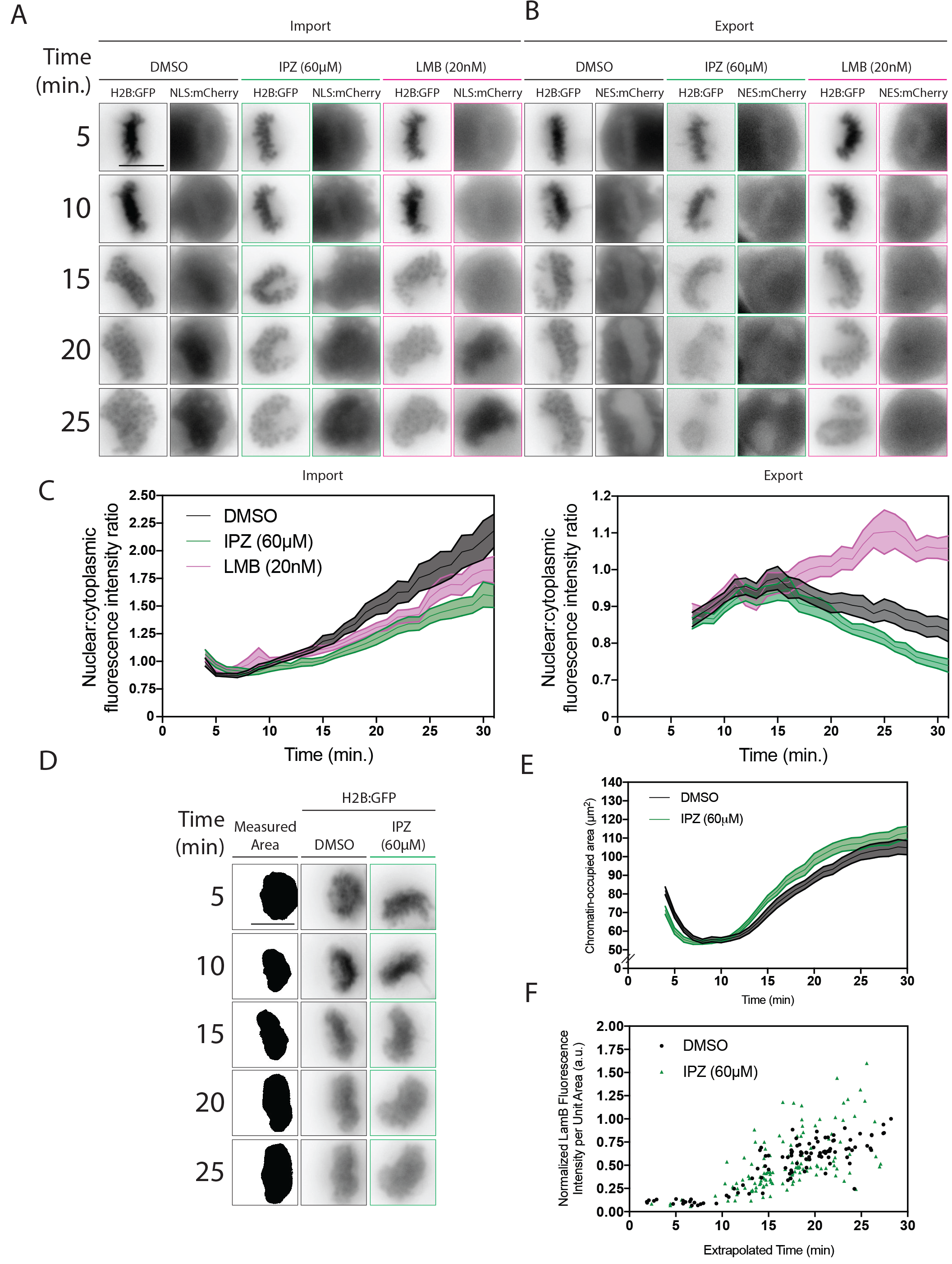
Nucleo-cytoplasmic trafficking inhibition at anaphase onset by IPZ and LMB. IPZ (60μM) and LMB (20nM) were used to target nuclear import and nuclear export respectively. **a,** Nuclear import was monitored with a nuclear import reporter (NLS:mCherry) during nuclear assembly. HeLa cells expressing H2B:GFP and NLS:mCherry were treated with DMSO, IPZ or LMB at anaphase onset. **b,** Similarly, nuclear export was monitored with a nuclear export reporter (NES:mCherry) during nuclear assembly. HeLa cells expressing H2B:GFP and NES:mCherry were treated with DMSO, IPZ or LMB at anaphase onset. **c,** NLS:mCherry and NES:mCherry fluorescence intensity was measured in the nucleoplasm and in the cytoplasm. **d,e,** Chromatin-occupied area was measured following IPZ treatment at anaphase onset in HeLa cells expressing H2B:GFP. F, Normalized LMNB1 fluorescence intensity in 3D maximum intensity projections at DNA. Extrapolated times were determined for individual nuclei based on chromatin-occupied area in Fig. S1E. Scale bars, 10 μm.

**Figure S2.**
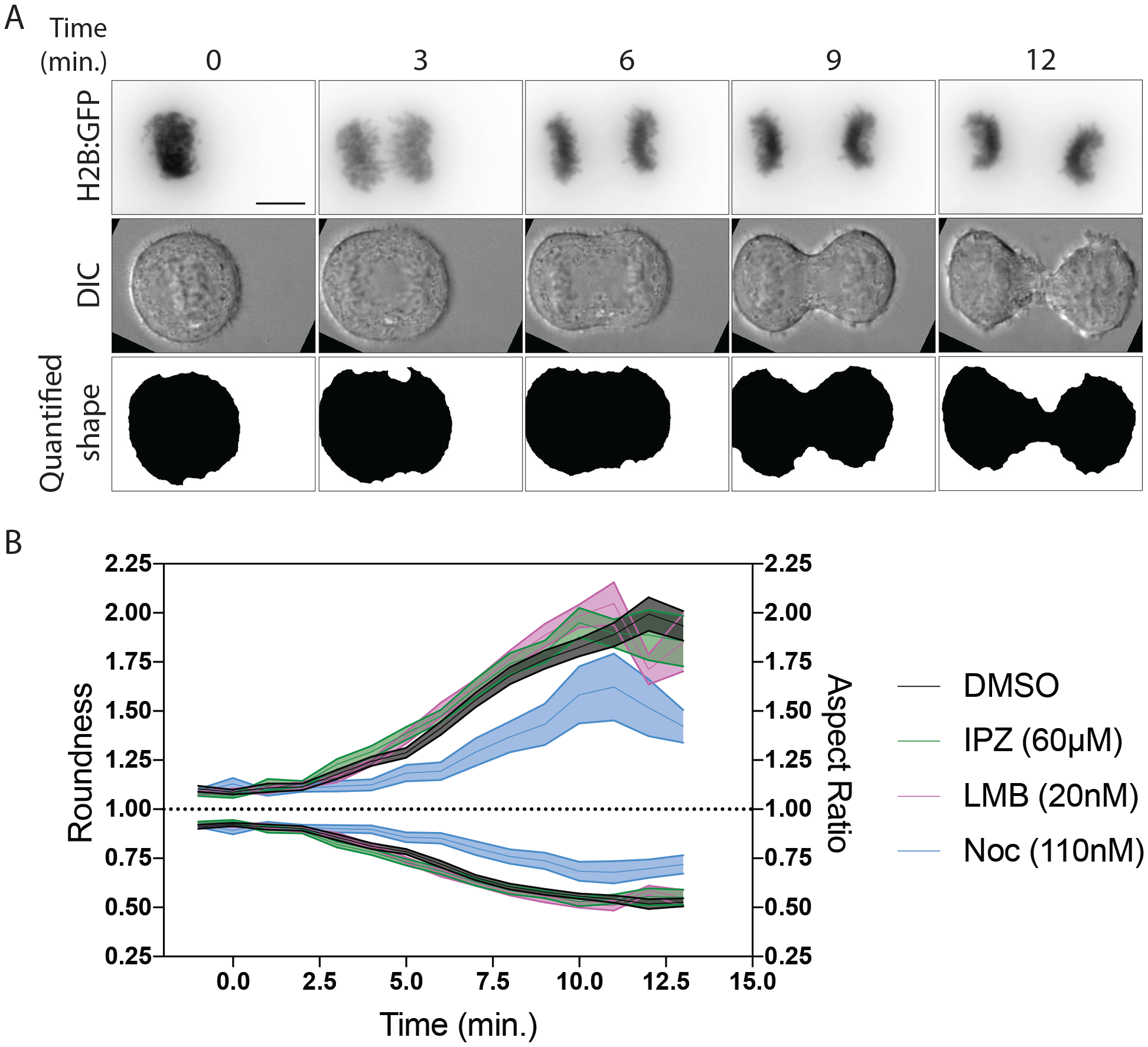
Treating cells with IPZ or LMB at anaphase onset does not affect global mitotic exit timing. **a,**HeLa cells expressing H2B:GFP were treated with DMSO or small molecule inhibitors at anaphase onset. Differential Interference Contrast (DIC) images were acquired in the middle slice of the acquired Z-stack and used for morphological analysis of mitotic exit progression. **b,** Morphological analysis and quantification of cell shape based on DIC images acquired in Fig. S2A was carried out using a custom ImageJ plugin. Nocodazole (110nM) slowed progression while IPZ and LMB had little to no effect on the timing of mitotic exit progression. Scale bar, 10 μm.

**Figure S3.**
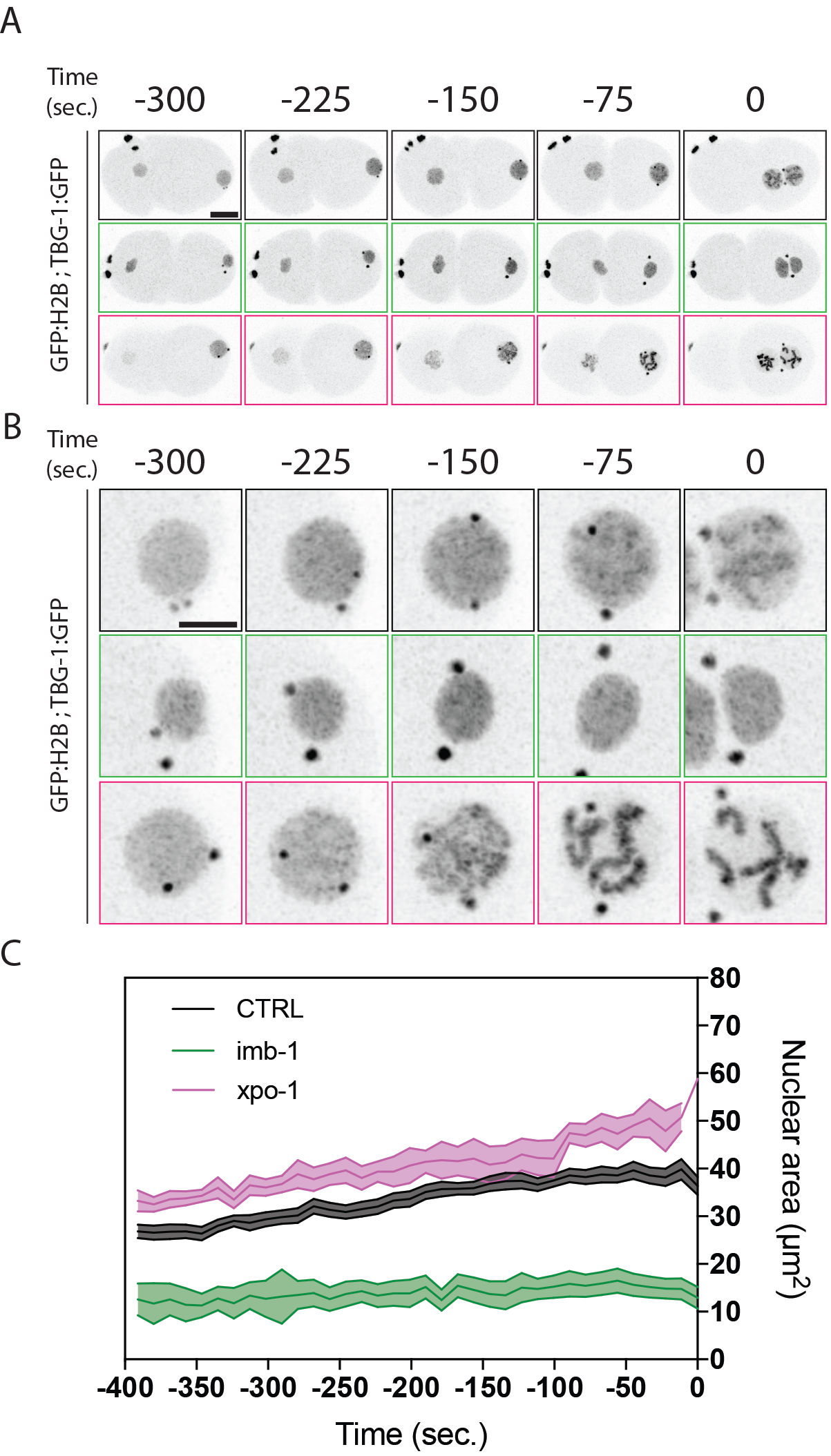
Targeting IMB-1 and XPO-1 via RNAi in the *C. elegans* embryo compromises nuclear expansion similarly to our mathematical model’s prediction. **a,** C. elegans L4 worms expressing GFP:H2B and TBG-1:GFP were depleted of target via dsRNA feeding. CTRL (L4440) worms were fed for 24 hours, IMB-1-targeted worms for 12 hours and XPO-1-targeted worms for 24 hours. **b,** Male pronuclei were imaged up to pronuclear meeting (t = 0 sec.) and c, pronuclear size was measured using a custom ImageJ plugin. Scale bar, 10 μm.

**Figure S4.**
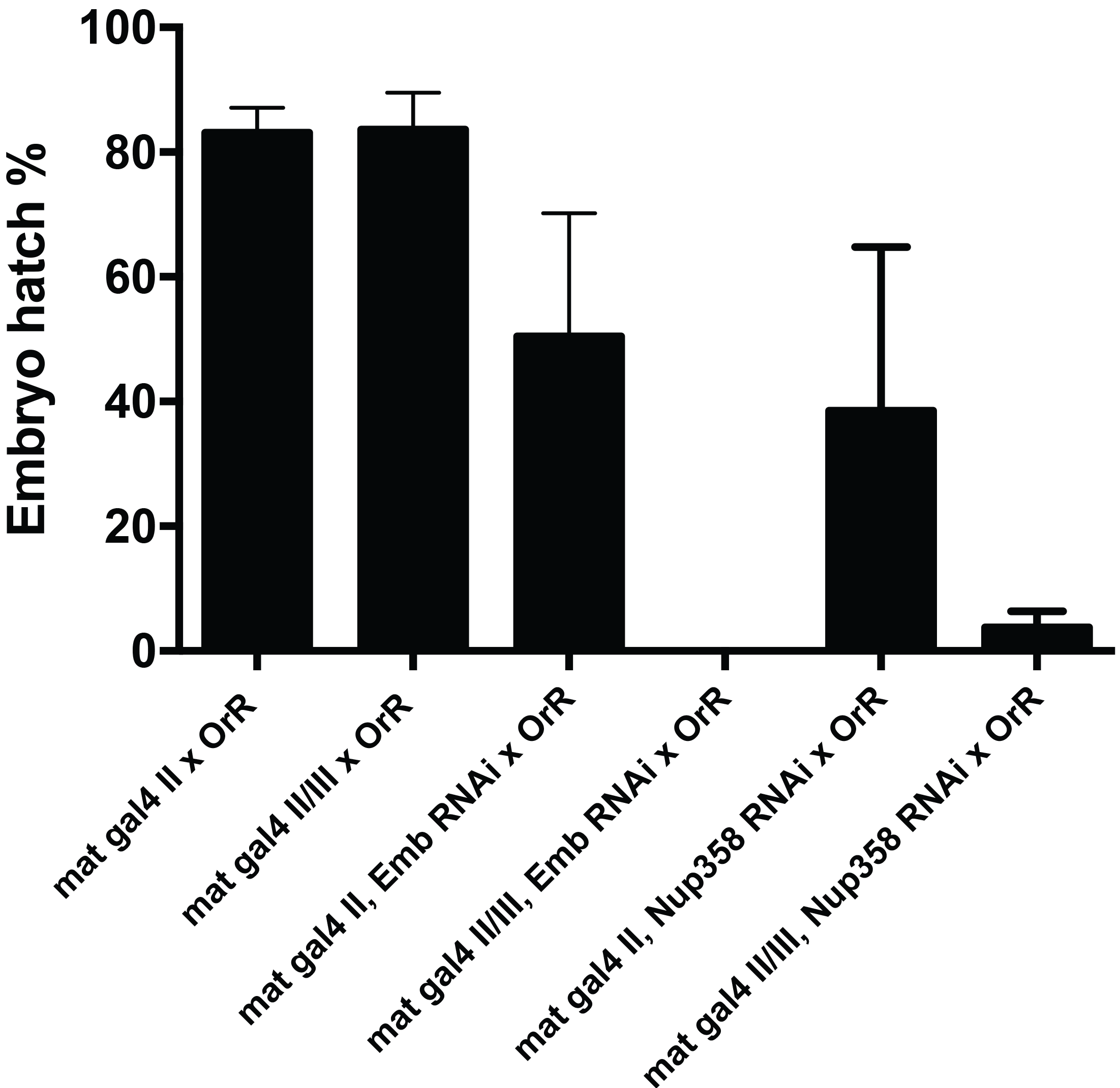
Depletion of Emb and Nup358 in the D. melanogaster embryo via RNAi leads to embryonic lethality. Embryos from mothers containing either one (mat gal4 II) or two (mat gal4 II/III) UAS-Gal4 drivers were used as controls. Mothers containing a single copy of either Emb-targeting UAS or Nup358-targeting UAS to drive dsRNA expression display increasing embryonic lethality when driven by a single copy of the mat gal4 driver and by two copies of the mat gal4 driver. Mothers expressing a single copy of the mat gal4 driver and of the UAS were used in Fig. 6. All mothers used were crossed with OrR (wild type) males.

